# Quantitative prey species detection in predator guts across multiple trophic levels by DNA shotgun sequencing

**DOI:** 10.1101/2021.04.01.438119

**Authors:** Débora P. Paula, Renata V. Timbó, Roberto C. Togawa, Alfried P. Vogler, David A. Andow

## Abstract

1. Metabarcoding has revolutionized the study of ecological communities, but PCR bias hampers quantitative analyses, as required in studies of trophic interactions. Direct DNA shotgun sequencing can avoid the amplification step when unassembled reads are mapped to a reference database, enabling identification and quantification of prey items.
2. Two feeding bioassays tested the precision and accuracy of quantitative assessments with the coccinellids *Harmonia axyridis* and *Hippodamia convergens*, the chrysopid *Chrysoperla externa*, and the aphid *Myzus persicae*. Guts were dissected and the total DNA extracted was directly sent to sequence by Illumina HiSeq2500 (insert 350 bp, PE 250). Predator gut content reads were blasted against an arthropod mitochondrial DNA reference database for taxonomic assignment and the matches curated for false positive prey identification using a series of bioinformatics pipelines, mostly in *R*. Taxonomic assignment through KrakenUnique, which is based on the counts of unique *k*-mers, was compared.
3. In a Prey Quantity bioassay, the number of prey reads was correlated to the amount of prey consumed and the elapsed time since consumption. In a Direct and Indirect Predation bioassay, prey was detectable in the predator 6 h after feeding (direct predation) and the prey’s prior prey was detectable 3 h after feeding (indirect predation). Detection of indirect predation was related to the species-specific decay rates of the two predators, but not to the order of predation.
4. We demonstrate that degraded prey DNA was quantifiable across trophic levels with high accuracy (98.4% positive predictive value) and taxonomic resolution.

## 1. Introduction

Species identification in environmental samples has been greatly enhanced by molecular methods. Several successful DNA-based methods rely on specific molecular markers, such as a single-locus or multiple barcodes in the nuclear (*e.g.*, ITS) or organellar (*e.g*., COI) DNA (Hebert et al. 2003; Hollingsworth et al. 2009) coupled with their enrichment by PCR. A key innovation has been metabarcoding (Valentini t al. 2009; Pompanon et al. 2014; De Barba et al. 2014), which uses “universal” primers to amplify and identify many taxa without the need to target specific species (Taberlet et al. 2012). This has led to hopes for, and controversies over the potential for quantification of community food webs and the population-level interaction strengths of prey and predators using relative read abundance (RRA) (Deagle et al. 2019). The main criticism of these metabarcoding enrichment methods is that PCR amplification bias, due to variation in primer efficiency across taxa and stochastic errors, may lead to species misidentification, lack of detection and over- or under-estimation of interaction strength (Yu et al. 2012; Deagle et al. 2014). Quantifying interaction strengths in ecological food webs remains crucial for understanding community dynamics (Pringle & Hutchinson 2020) and has significant implications for pest management and species conservation.

More recently, DNA-based species identification methods were developed that rely on the ‘natural’ enrichment associated with longer and more abundant target sequences, such as mitochondrial DNA, plastids, and nuclear ribosomal DNA clusters (Li et al. 2015). These include mito-metagenomics (Zhou et al. 2013; Tang et al. 2014; Crampton-Platt et al. 2015) and genome skimming (Linard et al. 2016), which generate assembled “superbarcodes” to increment reference databases. These sequences are a formidable resource for identification of any short fragments of DNA, including the minute amounts of digested prey DNA in predator gut contents. Thus, prey detection through the direct DNA shotgun sequencing (DDSS) of the predator gut contents has potential to be quantifiable, without DNA sample enrichment, opening up the possibility of quantitative interpretation of the data with high positive predictive value (Srivathsan et al. 2015; Paula et al. 2015; Paula et al. 2016). However, controlled experiments are needed to determine the feasibility of DDSS to quantify predator-prey interactions in food webs.

Here we conducted two feeding trials and developed an improved bioinformatics method to demonstrate that DDSS enables read numbers to be quantitatively interpreted as the number of prey items and the elapsed time since predation. To delineate the power and limitations, we estimate: how the amount of prey consumed and time after consumption affect prey detection; the likelihood of prey detection in relation to the number of prey consumed and time since consumption; the likelihood of false positive prey detections; the likelihood of distinguishing direct and indirect predation; and the positive predictive value, which is the percent of all positive identifications that are true positive detections. These findings also extend to the detection of prey items consumed indirectly by a predator (secondary predation). These findings validate the use of ‘PCR-free’ short-read data to identify species in environmental samples through mapping to reference genome databases (Anikeeva et al. 2019).

## 2. Material and Methods

### 2.1) Feeding bioassays

The predators were the ladybird beetles *Ha. axyridis* and *Hi. convergens* (Coleoptera: Coccinellidae), and the lacewing *C. externa* (Neuroptera: Chrysopidae); and the aphid prey *M. persicae* (Hemiptera: Aphididae). They were reared as presented in Supplementary Information. There were two bioassays:

#### a) Prey Quantity bioassay (Fig. 1a)

Twenty-four hour starved 3^rd^ instars of *Hi. convergens* were fed with zero (control-group), one, three or six apterous *M. persicae* to investigate if the number of prey reads detected was linearly related to the number of prey consumed. The bioassay was performed inside controlled environment chambers at 25°C, 16:8 h L:D. Predators were given 1 h to feed, and if they did not feed, they were discarded. Ten replicate predators (five males and five females) were sacrificed at each time point after feeding: 0 or immediately after feeding, 3, 6 and 9 h after feeding. At the given time point after feeding, predators were stored in 95% ethanol at −20°C.

**Fig. 1.**
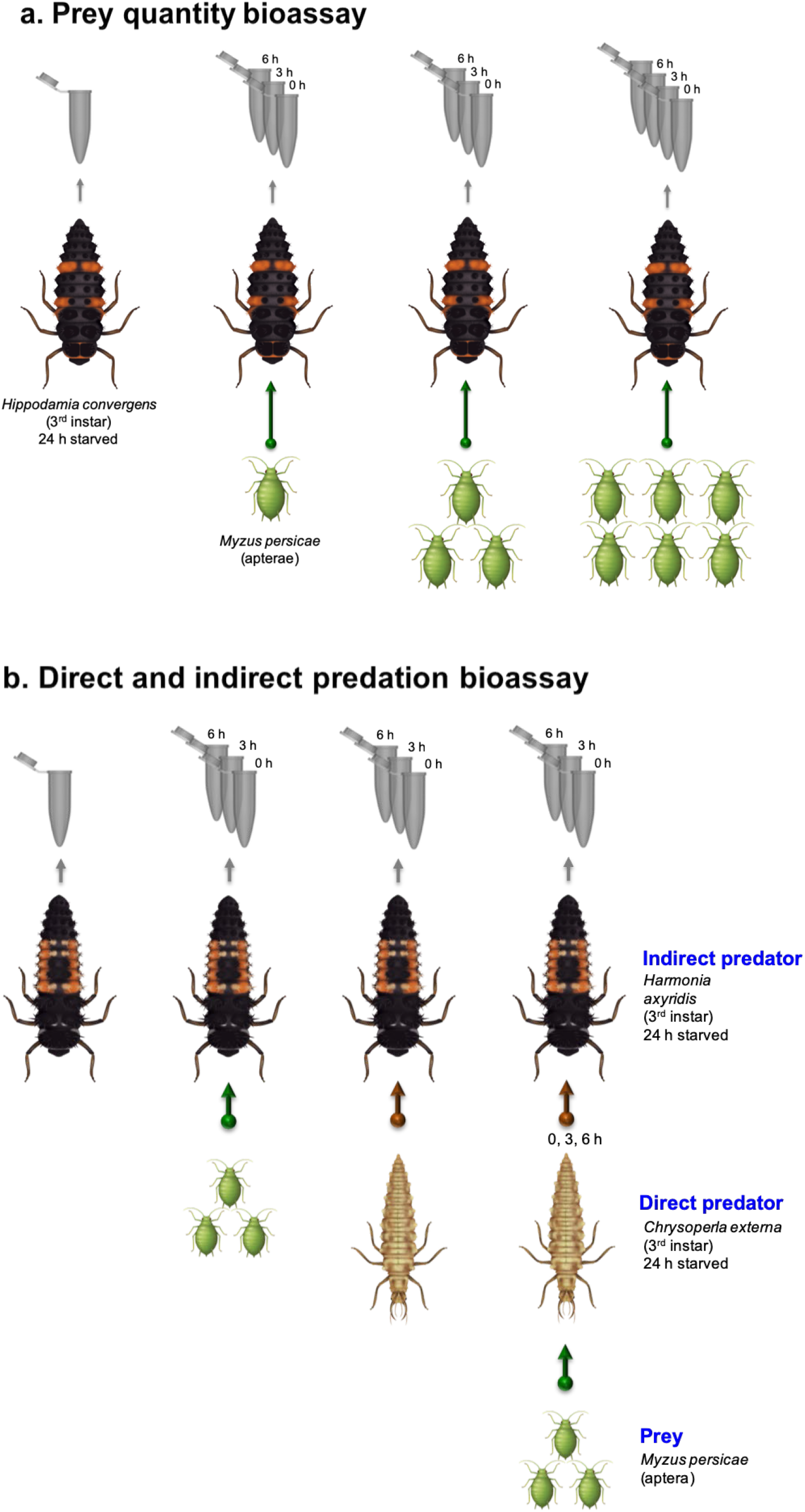
Feeding trial bioassays for prey detection using direct DNA shotgun sequencing (DDSS) of arthropod predator gut contents. Arrow directions indicate predation. Predators were offered no prey (control-group) or prey, and their gut contents were analyzed immediately after feeding or three or six hours after feeding. Insect drawings are copyrighted ©Carmen Martin 2018 and used with permission.

#### b) Direct and Indirect Predation bioassay (Fig. 1b)

Indirect predation (or secondary predation) is when a predator consumes a prey by consuming a predator that consumed the prey. Twenty-four hour starved 3^rd^ instars of *Ha. axyridis* were used as the top predator, *C. externa* was used as the intermediate predator and *M. persicae* was used as the prey. A 3^rd^ instar starved *C. externa* was provided zero (control) or three apterous *M. persicae*, and 0, 3 or 6 h after ingesting *M. persicae* prey, it was given to a 3^rd^ instar *Ha. axyridis*, which was held for 0, 3 or 6 h after ingestion of *C. externa* before storing in 95% ethanol at −20°C. We also included an unfed *Ha. axyridis* control and *Ha. axyridis* directly ingesting three apterous *M. persicae* and stored 0, 3, and 6 h after ingestion. Methods followed those of the Prey Quantity bioassay. The aim was to estimate the decay in detection of the aphid prey after direct versus indirect predation; the decay in detection of *C. externa* in *Ha. axyridis* irrespective of the previous feeding history of *C. externa* on the aphid prey; and the correlation of the detection of *M. persicae* and *C. externa* reads in *Ha. axyridis*.

### 2.2. Sample preparation for DNA sequencing

Whole gut contents of all of the predators were dissected under a microscope (30× magnification) using entomological scissors and forceps sterilized exclusively for each sample. Dissected gut content replicates from the same treatment/elapsed-time-of-consumption/bioassay were immediately pooled and immersed in microtubes on ice containing lysis buffer from the DNeasy Blood and Tissue (QIAGEN) kit. Total DNA extraction was performed using this kit according to manufacturer instructions. Total DNA yield was estimated by dsDNA High Sensitivity Assay in a Qubit 2.0 (Life Technologies). The amount of DNA was standardized across samples to 1 μg before drying in a Vacufuge Plus Vacuum Concentrator (Eppendorf™). Samples were used for TruSeq Nano library construction with 350 bp insert size (50-200 ng of DNA/sample). DNA library construction and sequencing were performed by the Roy J. Carver Biotechnology Center at the University of Illinois (USA) multiplexing all the libraries in one lane of Illumina HiSeq2500 (paired-end read, 250 bp).

### 2.3. Direct DNA Shotgun Sequencing (DDSS) analysis

The quality assessment for each library was done using FastQC (v.0.11.3) (Andrews 2010). Low quality sequences (Phred<30) and library index adaptors were trimmed by Fastqc-mcf (v.1.04.807, Aronesty 2011) and Cutadapt (v.1.9.1, Martin 2011). Retained good quality Fastq reads were converted to Fasta format by SeqTK (v1.2, Shen et al. 2016). The reference database was prepared by downloading from GenBank all the Arthropoda mitogenome sequences available at the time (April 2020) with 4071 species, which 35 were from the Coccinellidae family and 38 from the Aphididae family. Any other part of the genome, in addition to or instead, could be used as a reference database. Unassembled raw paired-end reads from each predator gut content were converted to Fasta format and aligned against the Arthropoda mitogenome reference database by BlastN v2.10.0+ (Altschul et al. 1997) using an E-value <1e-30, output format specifiers-outfmt “5 -> if ($3 > 90 && $4 > 100)”, dust=yes, word_size=15, and num_threads=20. The BlastN xml output result was converted to a tabular format and filtered for identity percentage above 90% and overlap length above 100 bp to reduce the size of the output file and because taxonomic resolution reduces poor at overlap lengths less than 100 bp (Meusnier et al. 2008). Each read can match multiple reference sequences and multiple locations of a single reference sequence (each is called a hit). Each hit match is a separate record. Prey detection was based on a threshold determination and a series of additional filtering steps to clean false positive detections, as summarized in Fig. 5 and presented in Supplementary Information. The thresholds for prey detection in our study could be determined as the data was from feeding trials, enabling us to distinguish between false or true prey species detections. However, for field samples, the prey species consumed are unknown, so one may consider using the thresholds determined in this work and to have a ‘predator-blank’ library made with the legs and/or antennae of the predator (i.e. a predator sample without any exogenous arthropod DNA) to assess potential false positive prey detection.

### 2.4. KrakenUniq analysis

Instead of conducting DDSS analysis, one may use a taxonomic sequence classifier, designed for metagenomics, that assigns taxonomic annotation to short reads based on the identification of unique *k*-mers for every species in a dataset. Based on Breitwieser et al. (2018), the reference database was reconstructed into a list of *k*-mers (default *k*=31) with associated taxonomic information (lowest common ancestor). Then it searched for *k*-mers and matched exactly query *k*-mers to the reconstructed reference database. Next, it counted the number of unique query *k*-mers and reads matching the taxa in the reference database and the coverage of the unique *k*-mers for each taxon in the database (workflow summarized in Fig. 5). To set a threshold number of unique *k*-mers to eliminate false positives, we established a threshold of 200 unique *k*-mers (average of 28 unique *k*-mers/read for non-predator species), which retained prey reads when rare, and optimized the tradeoff between retaining prey reads and eliminating false positive reads. Finally, the number of reads for each species in the control libraries (predator only, no prey consumed) was used as a blank to subtract from all of the experimental libraries.

### 2.5. Statistical methods

The statistical analysis follows that in Paula et al. (2015), and is summarized briefly here. Detection of a prey read is a stochastic process associated with sampling the predator gut, extracting DNA, and creating and sequencing the library. Assume that each read had the same probability of detection, the number of observed prey reads in a library will follow a Poisson process. We used Bayesian statistics to estimate a posterior distribution for the observed prey reads with the non-informative Jeffries prior distribution. The posterior distribution was normalized to the same initial DNA sample weight (200 ng) and the average library size after QC (2750640 reads). The mean, variance and 95% credibility interval of the ln transformed normalized posterior distribution were calculated from 200000 random draws from the distribution using a customized *R* script. Our previous work (Paula et al. 2015) indicated that prey read decay in a predator gut followed a first order exponential decay process, which means that decay occurred at a constant rate. Therefore, we estimated the decay rate (*d*), the ln initial number of prey reads (ln(*n_0_*)) and their standard errors from a weighted linear regression of time after prey ingestion on the estimated ln prey read number with inverse variance weights using glm in the lme4 package in *R*. We estimated the maximum detectability period (*D_max_*) and its variance and 95% credibility interval by Monte Carlo simulation (Paula et al. 2015). Four sets of regressions were conducted: effect of prey number; decay of *M. persicae* in *Ha. axyridis*; decay of *M. persicae* reads in *C. externa* larva; decay of *C. externa* reads in *Ha. axyridis*. They are described in Supplementary Information. All of the regressions and tests were conducted for the six best thresholds to determine the sensitivity of the result to the threshold using glm in the lme4 package in *R*.

## 3. Results and Discussion

### 3.1. Number of prey quantitatively affects the number of prey reads detected and the maximum detectability period

In the ‘Prey Quantity’ bioassay (Fig. 1a), true prey species were detected in all libraries except in control libraries, and when only one prey specimen was consumed ≥ 6 h ago. All of the detectability parameters (initial number of reads *n*_0_, decay rate *d*, maximum detectability period *D_max_*) were within the range that had been previously reported for detection by species-specific PCR of equivalent prey types and stages (Greenstone et al. 2014). The number of prey reads detected after filtering immediately after feeding ranged between 174 and 2814, which was normalized to an equivalent amount of DNA in the original sample and average library depth after quality check (QC). The normalized *n_0_* was significantly higher when a higher number of prey were consumed (*p* = 6.475E-06), which was found by others using qPCR (Weber & Lundgren 2009). The decay rate of *Myzus persicae* reads (aphid prey) fitted a linear decay process (Fig. 2a) and was not influenced by the number of prey consumed (rate ranged from −0.98 to −1.02 h^−1^, *p* = 0.9913), as expected (Hoogendoorn & Heimpel 2003; Deagle et al. 2006; Greenstone et al. 2007; Gagnon et al. 2011). Considering that detectability half-life is −1/*d*, these decay rates mean that the number of prey reads decreased by half every 1 h of predator digestion. The higher the number of prey consumed, the longer was the *D_max_* (7.61 to 11.82 h, *p* = 0.0058).

**Fig. 2.**
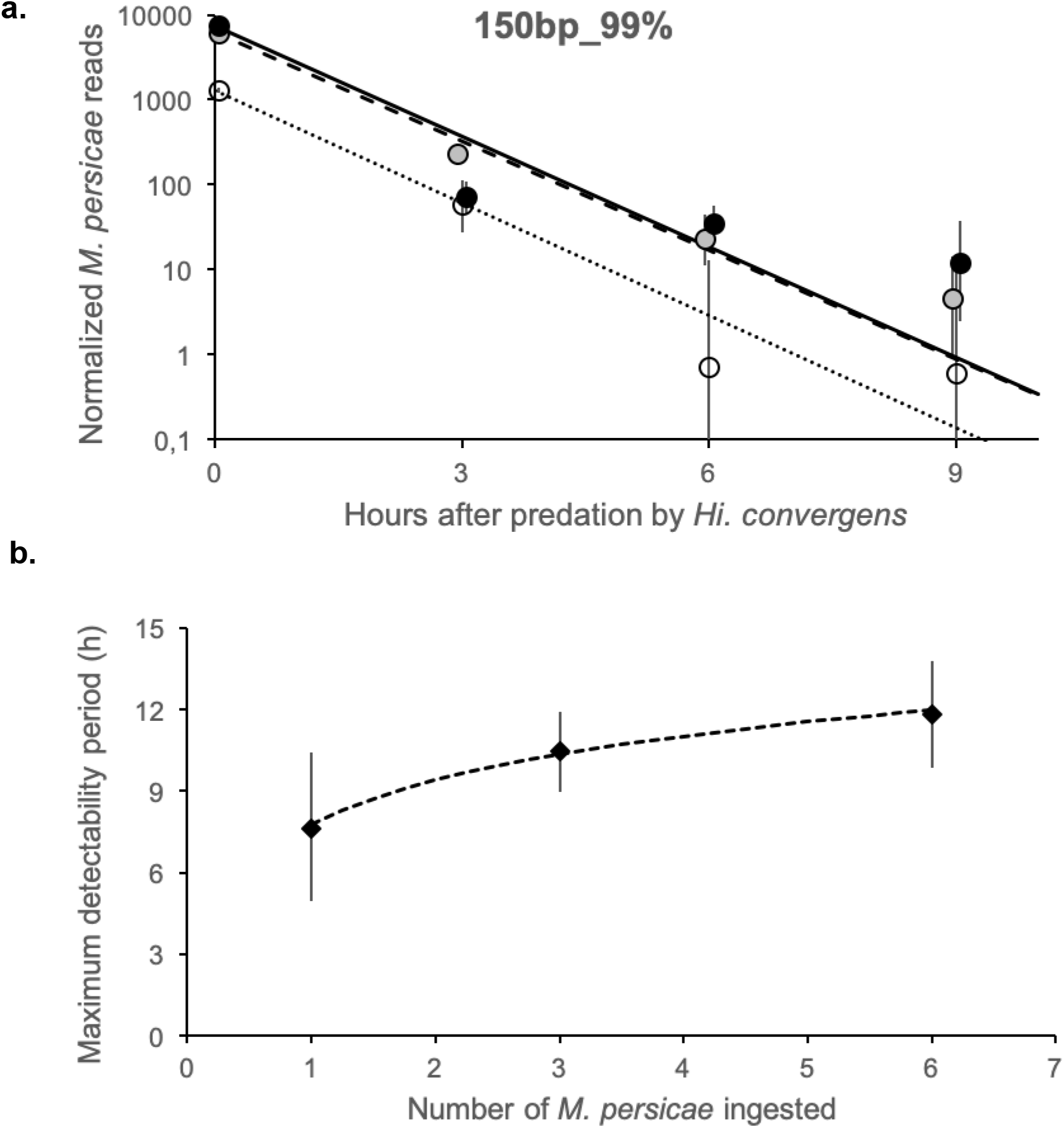
*Myzus persicae* read detectability in the gut of the predator *Hippodamia convergens* related to different numbers of prey consumed. **a**, Decay and **b**, Maximum detectability period (*D_max_*). Observed data with 95% CIs and weighted linear regression lines.

We predicted that the initial number of reads (*n*_0_) detected would be directly proportional to the number of prey ingested, if read detection were quantitatively related to the number of prey consumed. In other words, ln(*n*_0_) ∝ *m*_1_ ln(*Mp*), where *Mp* is the number of *M. persicae* prey reads detected and the slope, *m*_1_ = 1. In addition, as the read decay rate, *d*, was constant over time, this predicts that ln(*n*(*T*)) ∝ *dT* after ingestion, where *n* is the number of reads detected at time *T* after ingestion. Finally, both predictions imply that *D_max_* ∝ ln(*Mp*). The multiple regression model based on the first two predictions, ln(*n*(*T*, *Mp*)) ~ ln(*n*_0_)+*m*_1_ ln(*Mp*)+*dT*, fit the data extremely well (*r*^2^ = 0.932) and the slope, *m*_1_, was not significantly different from 1 (Supplementary Table S1, *t*_9_ = 0.5009, *p* = 0.628). The regression for the third prediction with *D_max_* also fit well (*r*^2^ = 0.988, Fig. 2b). These results indicate that the number of prey reads detected was quantitatively related to the number of prey consumed and the time since consumption, and that the detectability period increases quantitatively with the number of prey consumed.

### 3.2. Indirect predation is detectable

In the ‘Direct and Indirect Predation’ bioassay (Fig. 1b), indirect predation on the aphid prey (*M. persicae*) by the primary predator, *Chrysoperla externa*, was detectable up to 7 h, which included 3 h after the aphids were consumed initially and digestion for 0 to 6 h in the secondary predator, *Harmonia axyridis* (*D_max_*, Supplementary Table S2). As expected, this was significantly shorter than direct predation on the aphid prey by *Ha. axyridis* (14 h). The detection of indirect predation on the aphid prey by DDSS was longer than that found by others using monoclonal antibodies (Harwood et al. 2001) (0 h) and similar to that using PCR (8 h, Sheppard t al. 2005). The normalized number of *M. persicae* reads detected immediately after being secondarily consumed by *Ha. axyridis* was 100, 86 and 50 for 0, 3 and 6 h after primarily consumption by *C. externa,* respectively. This was significantly lower than the 2316, 1090 and 76 normalized *M. persicae* reads detected 0, 3 and 6 h after direct consumption by *Ha. axyridis* (Fig. 3a, *p* = 0.00008, Supplementary Table S3). However, there were no significant differences in the decay rate of *M. persicae* reads in the *Ha. axyridis* after being directly or indirectly consumed (*d*_indirect predation_ = −0.11 to −1.00 h^−1^; *d*_direct predation_ = − 0.32 h^−1^, *p* = 0.2627, Fig. 3b, Supplementary Table S4). Also, there were no significant differences in the decay rate of the aphid in *C. externa* controlling for the time *C. externa* was consumed by *Ha. axyridis* (*p* = 0.6328, Fig. 3c, Supplementary Table S5). Although the decay rate of *M. persicae* reads in *Ha. axyridis* versus *C. externa* was not statistically different (*p* = 0.0965, Supplementary Table S6), it tended to be faster in the *Ha. axyridis* (*d_H. axyridis_* = − 0.48 to −1.00 h^−1^; *d_C. externa_* = −0.11 to −0.62 h^−1^, Fig. 3d). Direct and indirect predation on *M. persicae* was not distinguishable.

**Fig. 3.**
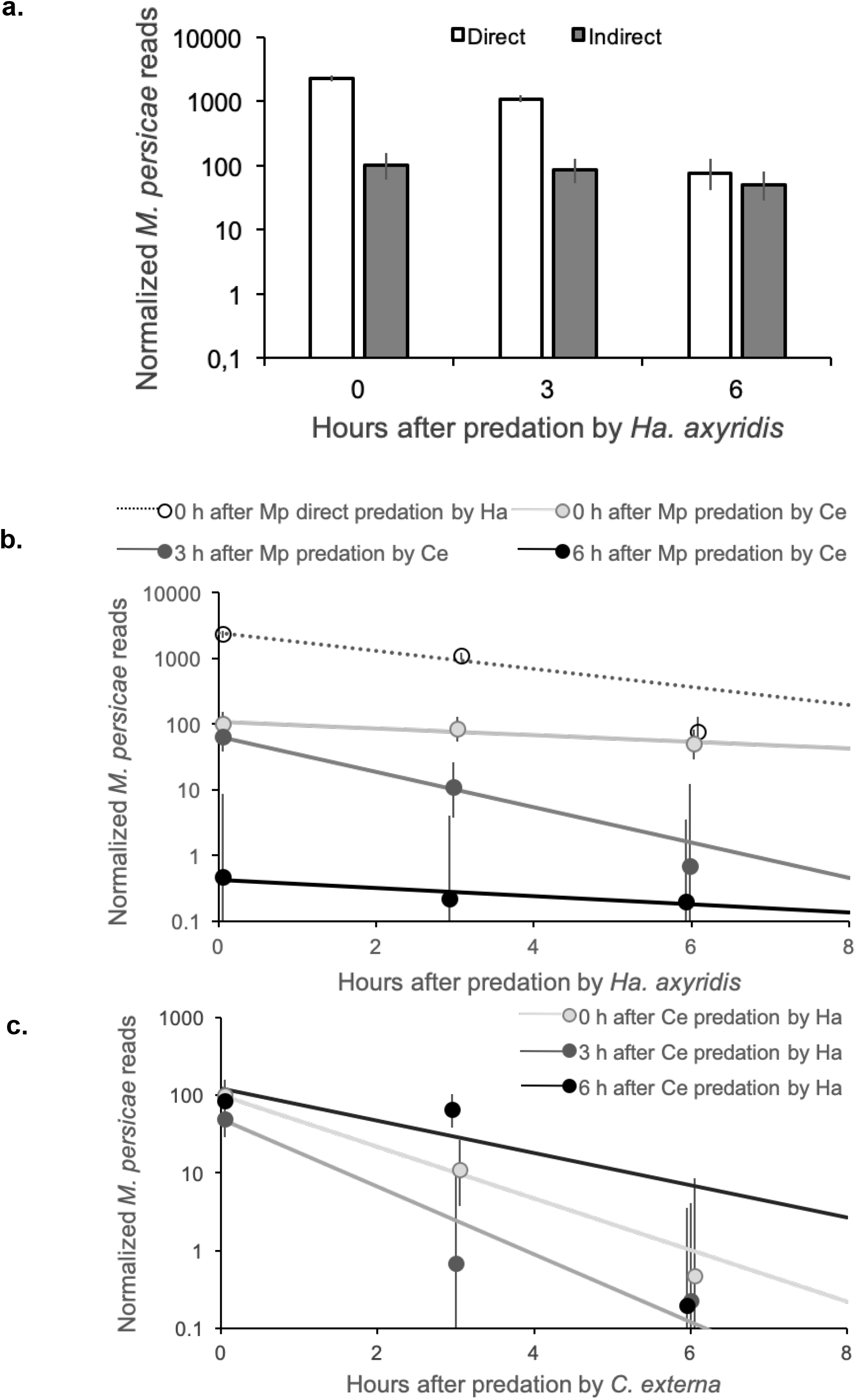
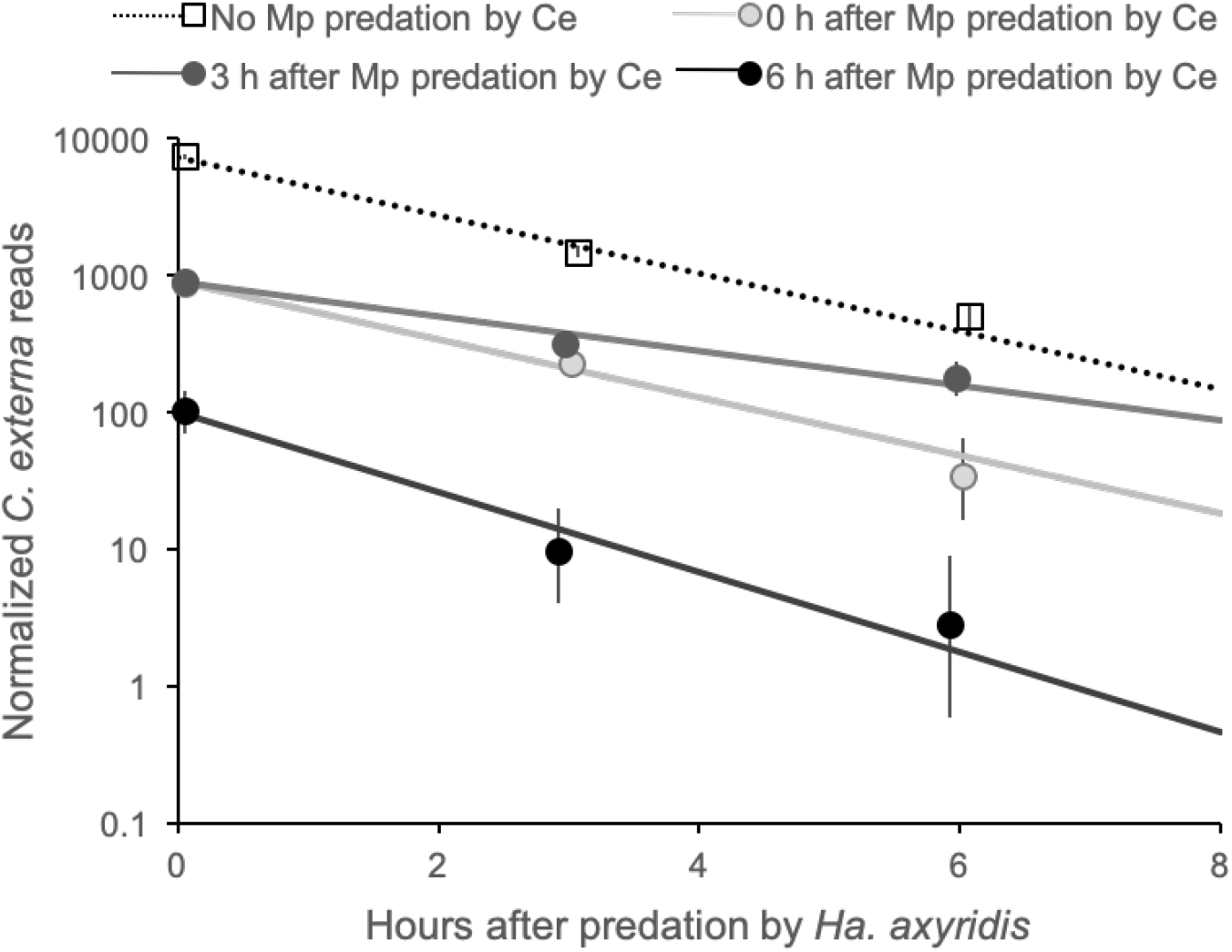
Prey detectability in intraguild and extraguild predation. **a**, Normalized number of *Myzus persicae* (Mp) reads detected in *Harmonia axyridis* (Ha) larvae after direct or indirect predation with 95% CIs. Direct: Mp was ingested by Ha; Indirect: Mp was ingested by *Chrysoperla externa* (Ce), which was immediately ingested by Ha. **b**, Decay of the detection of 3 Mp in Ha after direct consumption or indirect consumption of Ce, which had consumed Mp. **c**, Decay of the detection of 3 Mp in Ce after Ce was consumed by Ha. **d**, Decay of the detection of Ce that consumed 0 or 3 Mp at 0, 3 or 6 h before being consumed by Ha.

If read detection of the aphid prey was quantitatively and independently related to the time in *C. externa* and *Ha. axyridis*, read detection would fit the multiple regression model, ln(*n*(*T_H.axyridis_*, *T_C.externa_*)) ~ ln(*n*_0_)+*d_H.axyridis_T_H.axyridis_* +*d_C.externa_T_C.externa_*. Decay rates of *M. persicae* prey directly or indirectly consumed were statistically independent (*F*_1,5_ = 0.0069, *p* = 0.9368), and the model fit the data extremely well (*r*^2^ = 0.906). This implies that detection of indirect predation was related to the species-specific decay rates of the two predators and not on which of the predators directly or indirectly consumed the prey.

Direct predation on *C. externa* by *Ha. axyridis* was detected in all of the libraries with *C. externa*. The initial normalized number of *C. externa* reads detected in *Ha. axyridis* varied from 102 to 7370, which is similar to the number of aphid reads detected in *Ha. axyridis* after direct consumption (2316 reads). Decay rates were similar for all *C. externa* feeding histories, indicating that previous feeding by *C. externa* did not affect its decay rate in *Ha. axyridis* (*p* = 0.1313, Fig. 4, Supplementary Table S7).

**Fig. 4.**
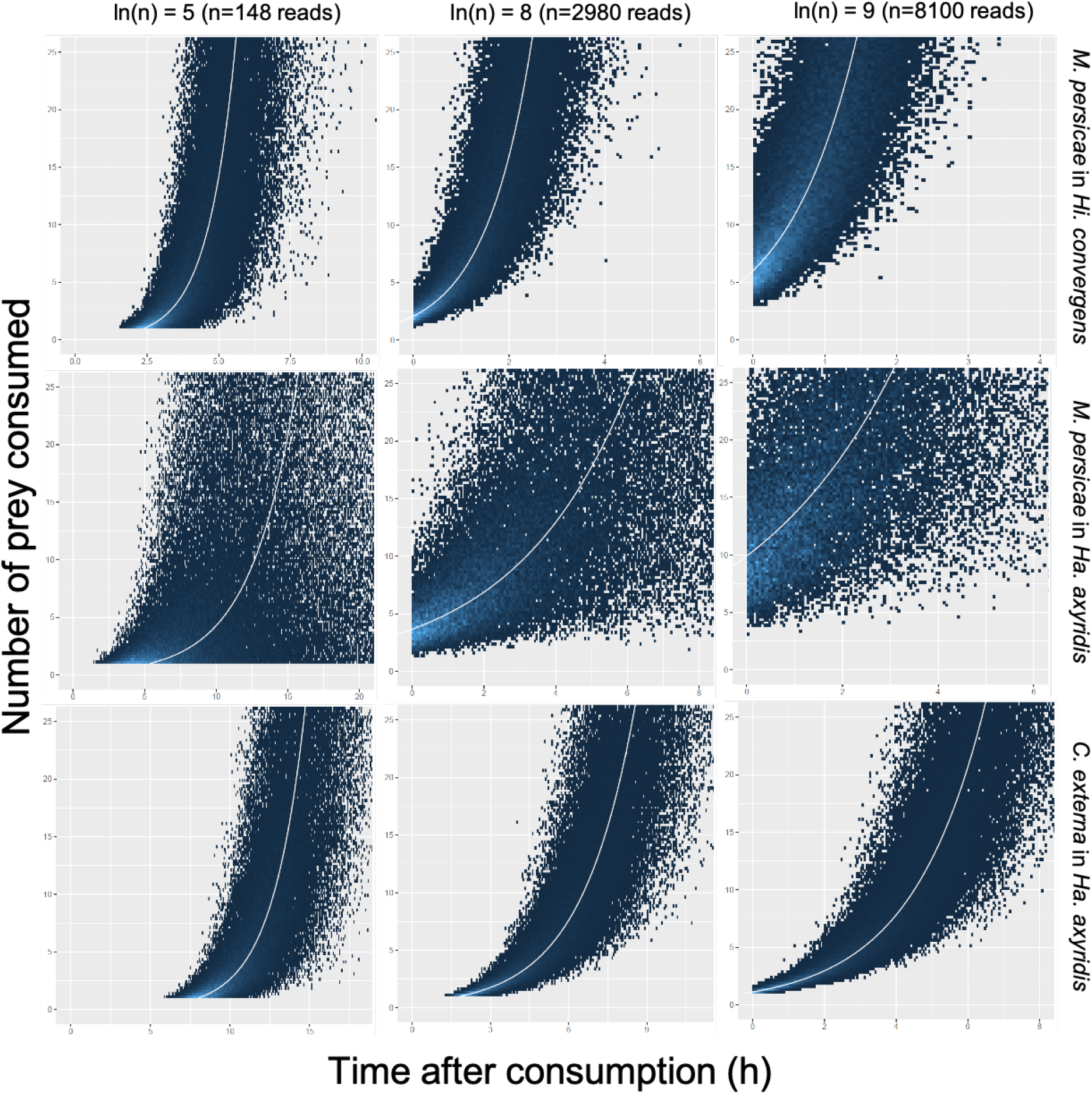
Heat maps of predicted number of prey consumed and time after consumption when detecting a hypothetical number of prey reads (columns: ln(*n*), *n* = number of prey reads) in a predator gut for *Myzus persicae* prey in the predators *Hippodamia convergens* and *Harmonia axyridis*, and *Chrysoperla externa* prey in *Ha. axyridis*. From 200000 Monte Carlo simulations of the inverse regression of our data, assuming normal parameters and prey reads are directly proportional to the number of prey consumed. White line is the deterministic prediction, i.e. it shows how the prediction would behave without parameter uncertainty; blue dots show prediction with uncertainty. Lighter blue regions have higher probability, which correspond to the prediction regions less influenced by parameter uncertainty.

**Fig. 5.**
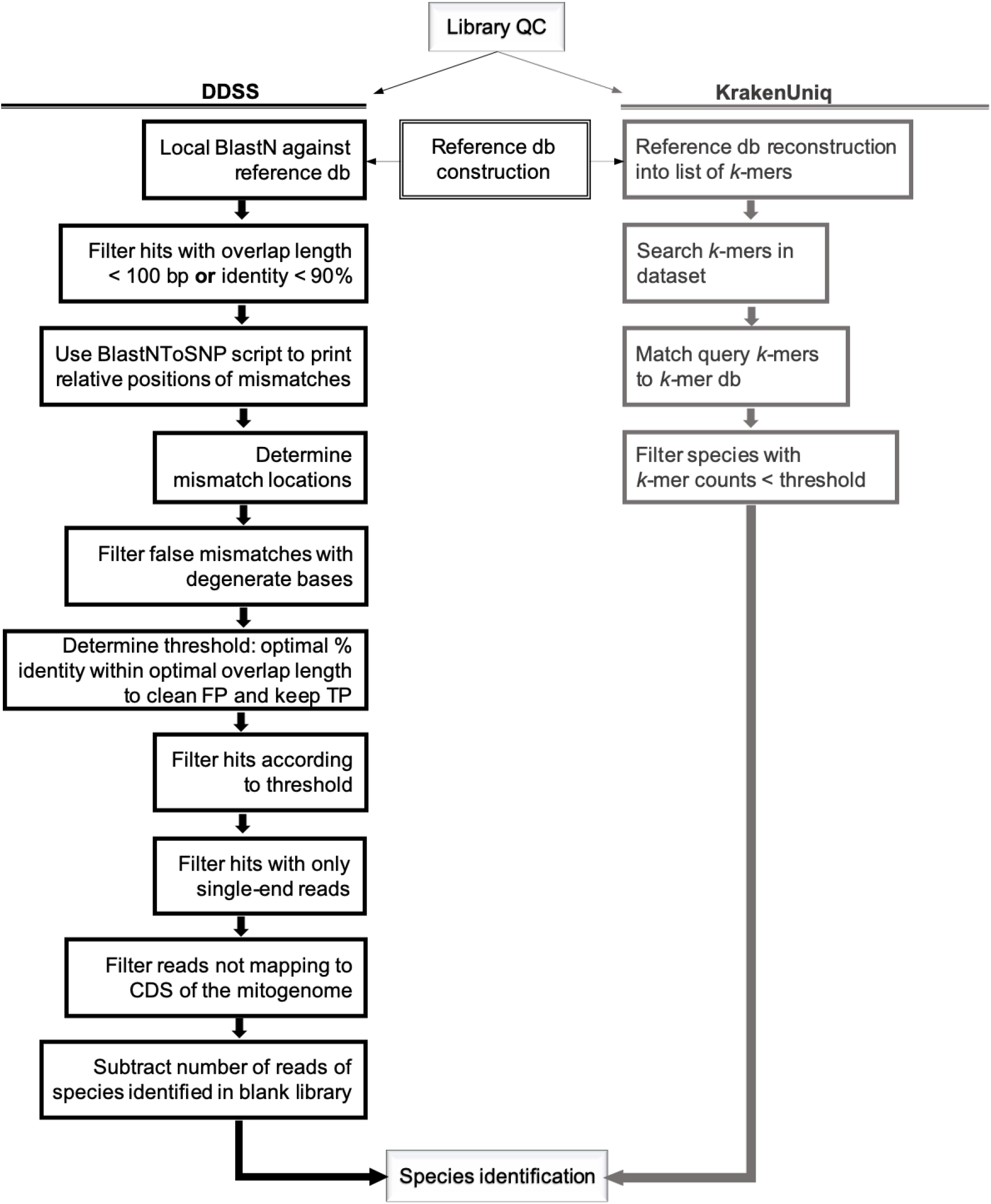
Bioinformatics workflows for arthropod predator gut content analysis to identify consumed prey species by improved Direct DNA shotgun sequencing (DDSS) or KrakenUniq (Breitwieser et al. 2018). QC: quality check; db: database; FP: false positive; TP: true positive; CDS: coding sequence.

### 3.3. Prey quantification is predictable

The direct relation between number of prey consumed, the time since consumption and the number of reads detected makes it possible to predict for predator samples from the field, when prey decay rates are known, how many prey were consumed how long ago using an inversion regression (Fig. 4). For example, in Fig. 4, a sample of the predator *Hippodamia convergens* or *Ha. axyridis* with the detection of 148 reads of *M. persicae* would correspond to the consumption of ca one *M. persicae* ca 2.5 h and 5 h after *Hi. convergens* and *Ha. axyridis* predation, respectively. It could also correspond to the consumption of 5 *M. persicae* ca 4 or 11 h after *Hi. convergens* or *Ha. axyridis* predation. As the probability is higher in the lighter blue region of the inversion regression graphs, which corresponds to a region less influenced by the uncertainty in the parameter estimates, the first interpretation is more likely than the second. Using the same rational, *n*= 148 reads of *C. externa* in the gut of *Ha. axyridis* would correspond to the consumption of ca one *C. externa* ca almost 8 h before *Ha. axyridis* was sampled. As prey read decay rate may be specific to every predator-prey species interaction, the decay rate must be determined before the number of prey consumed and time since consumption can be predicted. Once determined, the decay rate could be used as reference for new studies, as indicated here for *M. persicae* consumption by *Hi. convergens* or *Ha. axyridis*, and *C. externa* consumption by *Ha. axyridis*.

### 3.4. Elimination of false positives

Despite the lack of prey DNA enrichment by PCR, which may generate false positives through amplification bias or contamination, some false positive identifications remained (Fig. 6). However, the positive predictive value (PPV), which is equal to 100*TP/(TP+FP), where TP=number of true positive prey reads and FP=number of false positive non-predator and non-prey reads, was 98.4%, indicating that the false positive reads were a small proportion of the total. In DDSS, errors of read counts arise due to the ambiguity of nucleotide matches between reads and reference sequence, which depend on the read quality and read length, the genetic distance of prey species and reference across conserved and variable gene regions, and the completeness and accuracy of the reference database. We evaluated 25 thresholds of overlap length and percent identity and determined that the threshold with overlap length of 150 bp and identity of 99% was the best at eliminating false positives and retaining true positive prey species (Supplementary Table S8 and Fig. S1-S4). The false positive species initially identified are listed in Supplementary Table S9 and they were categorized into three types: potential cross-contaminants; mismatch species; and other species. ‘Potential cross-contaminants’ were species that were species used in the bioassays or other bioassays conducted at the same time. ‘Mismatch species’ were species in the same taxonomic family as one of the species in the experiment. The third group are ‘other species’, which were unrelated to any species in the experiment.

**Fig. 6.**
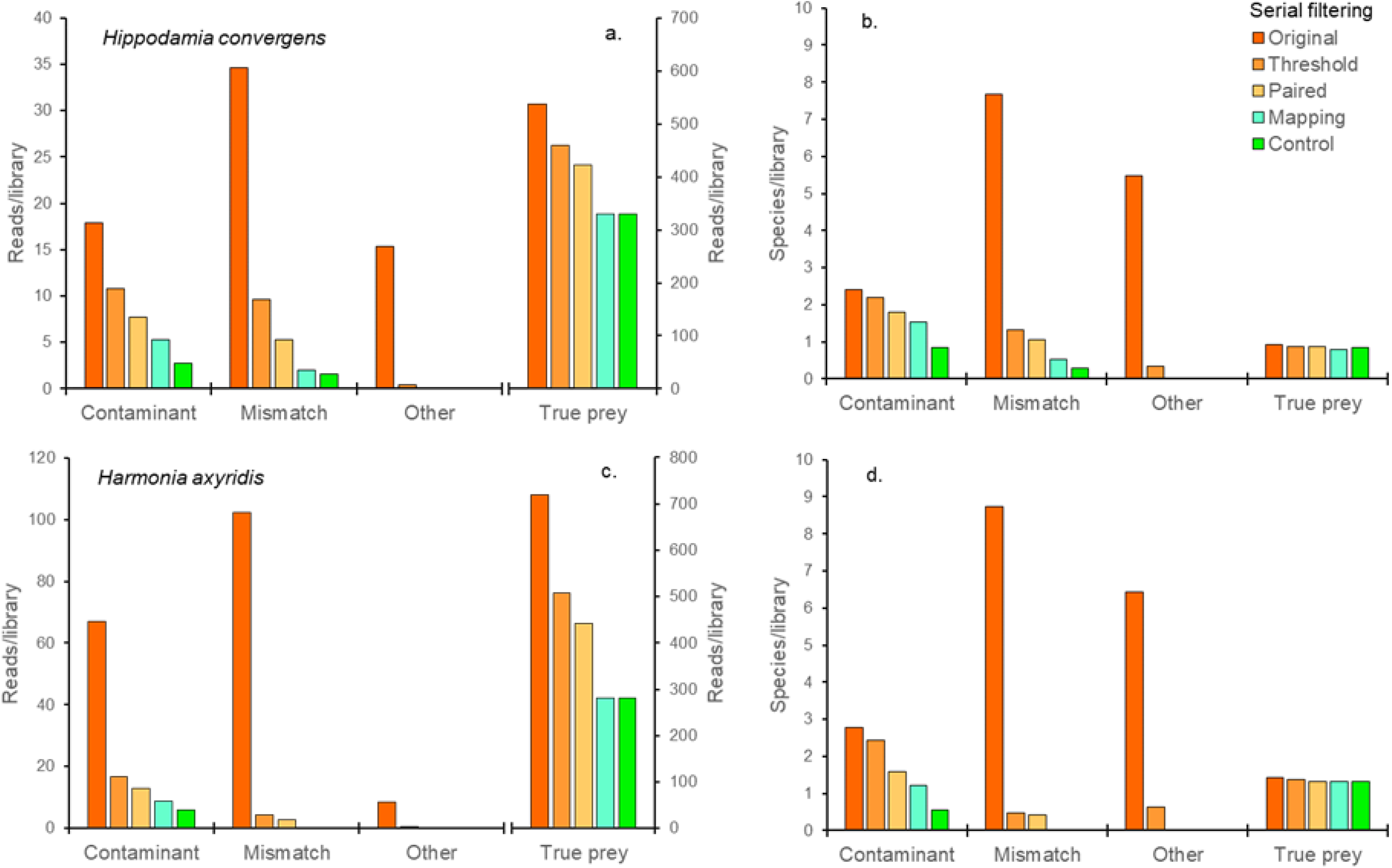
Serial filtering (indicated by different colored bars) of the average number of reads and species per library for three kinds of false positive detections (potential cross-contaminants, mismatch, and other) and detections of the true positive target prey. The four serial filters are: (1) Threshold: removal of reads below the minimum overlap length (150 bp) and percent identity (99%); (2) Pairing: removal of reads that are not paired forward-reverse reads with the same hit species; (3) Mapping: removal of reads that do not map to coding regions of the reference mitogenome; (4) Control: subtraction of reads in the blank library (no-prey consumed by the predator). a and b are for libraries with *Hippodamia convergens* as predator (Prey quantity bioassay) and c and d are for libraries with *Harmonia axyridis* as predator (Direct and Indirect Predation bioassay). a and c show how reads are filtered, and b and d how species detections are filtered. The false positive detections are: potential cross-contaminants, where DNA of species used in the bioassays leaked between libraries; mismatch species, which are in the same taxonomic family as n species used in the bioassays; and unrelated species (referred as other). The collateral effect of filtering on the detection of true prey reads is also shown (True prey).

Filtering reduced the number of false positive reads and species (Fig. 6). ‘Other species’ were almost completely eliminated with the threshold of overlap length of 150 bp and identity of 99%, and were completely eliminated when filtering for paired end reads with the same species identification. ‘Mismatch species’ were greatly reduced at the 150 bp/99% identity threshold and by only matching to coding regions of their reference genome. Mismatches may arise in part because they match a more conserved region of the mitogenome, which may not discriminate against the reference genome of the experimental species. Finally, ‘potential cross-contaminant’ species were probably a mixture of mismatches and contaminants, as they are all related to an experimental species similar to ‘mismatch species’. Reads of these species were considerably reduced by the 150 bp/99% identity threshold, similar to but not as much as for ‘mismatch species’. The subtraction of the control reduced these species and reads the most of all of the filtering steps (about 50%), which is consistent with them being contaminants. In short, the filtering method eliminated all reads and species of the other species (100% PPV), and only 13% of the remaining false positive reads were mismatch species (99.8% PPV).

While a low number of reads has been used to eliminate false positives in metabarcoding, this is inappropriate for DDSS. Because prey DNA is not amplified, true positive prey detections can have a low number of reads. Therefore, using read numbers to clean remaining false positives may also remove some true positives, especially of rare prey or prey consumed a long time ago. We suggest that most, if not all, remaining false positives can be eliminated by having replicate samples and using site occupancy models to distinguish true and false positive determinations after bioinformatics filtering. For example, we had three replicate libraries with one *M. persicae* prey consumed by *Hi. convergens*, and while all three libraries detected *M. persicae*, none of the five remaining false positive species were found in more than one of the replicate libraries.

To test if the false positive rates could be reduced using a taxonomic classification method other than sequence similarity with Blast, the datasets after QC were analyzed by KrakenUniq (Breitwieser et al. 2018), a classifier based on counts of unique *k*-mers. KrakenUniq detected a similar number of prey and predator reads as the DDSS workflow (Tables 1 and S10). It detected true prey species in one library (five reads) where none were detected with the DDSS workflow, however, it also detected more false positives than the DDSS workflow with a PPV of only 90.5% (~10% of detections were false positives). In addition, it retained a detection of an ‘other species’, while none of these were detected with the DDSS workflow.

**Table 1.** Number of reads of true positive species (prey) and false positive species (potential cross-contaminant, mismatch and other) and positive predictive value (PPV) for KrakenUniq and DDSS.

|  | Prey | Potential cross-contamination | Mismatch | Other | PPV (%) |
| --- | --- | --- | --- | --- | --- |
| KrakenUniq | 12222 | 1195 | 25 | 58 | 90.5 |
| DDSS | 10286 | 148 | 22 | 0 | 98.4 |

Undoubtedly, the biggest challenge for all environmental DNA methods that identify species is to discriminate between false positives, i.e. detection of species that were not in the sample, and true positives, the species that are actually present in the sample. In metabarcoding, this has been handled using several bioinformatics methods, including Perseus to remove chimeras after assembling paired-end reads (Quince et al. 2011), Obiclean to identify and eliminate sequencing errors (Shehzad et al. 2012; Boyer et al. 2016) and cluster-free filtering based on control samples to established non-arbitrary cleaning thresholds (Corse et al. 2017). However, typically more arbitrary methods are required to remove false positive species, including filtering by minimum amplicon sequence count and length (Reeder & Knight 2010; Quéméré et al. 2013; De Barba et al. 2014), removing singleton reads (Yu et al. 2012) and removing species that are not known to be present in the sampled environment (De Barba et al. 2014). When there are multiple replicate samples, ecological site occupancy models have been used to estimate detection probabilities for each species, which can then be used to filter species (Royle 2006; Miller et al. 2011; Schmidt et al. 2013; Ficetola et al. 2016; Lahoz-Monfort et al. 2016) using some detection probability threshold. Some of these methods can be applied to DDSS.

### 3.5. Benefits and limitations

The primary benefit of using a method without DNA enrichment, such as DDSS, is the preservation of the original prey DNA composition, which enabled quantitative interpretation of the data. Although the number of reads in metabarcoding studies is sometimes interpreted quantitatively, several studies have shown that this is not yet robust (Piñol et al. 2014; Elbrecht & Leese 2015). As demonstrated in this study and in Paula et al. (2015), the fact that in DDSS prey reads decay at a constant rate indicates that the number of reads detected is quantitatively related to the elapsed-time since predation. Moreover, the initial number of prey reads detected was directly related to the number of prey consumed and the decay rate did not depend on the number of prey consumed. Another advantage is that the type of reference database (genes from organellar DNA, rRNA gene cluster, etc.) can be chosen after the acquisition of the data and with virtually unconstrained possibilities. This allows for the detection of unanticipated exogenous DNA originating from different interactions of the predator, for example detection of parasites and symbionts associated with the predator or even symbionts of the prey (Paula et al. 2015; 2016). The only limitation to detection is the availability of genome sequences, but as thousands of well assembled genomes are becoming available (Earth Biogenome Project reference here), the information content of the shotgun mixture is constantly increasing.

There is no particular reason to suspect that the sequences in a potential reference database used in DDSS (e.g., mitogenome, plastid, nuclear ribosomal DNA) would have higher or lower accuracy than the shorter barcode sequences used in metabarcoding (e.g., Folmer region of COI). However, the comprehensiveness and redundancy of potential DDSS reference databases are currently lower. For example, there are over 1.7 million BOLD accessions for COI compared to 14719 GenBank accessions for insect mitogenomes (August 2020). Therefore, at low threshold values and given a taxonomically diverse prey community, there should be a higher likelihood of detecting true positive prey species targeting the COI gene. However, as the database of mitogenome sequences increases rapidly, this difference will decrease over time and the greater sequence information from the full mitogenome increases the detection limit by ~20× over the short COI fragment.

A potential limitation of the DDSS approach is the cost of data acquisition. DDSS requires that every sample has an individual library to preserve the sample identity because a sample tagging method has not yet been developed that allows different samples to be multiplexed. However, as the research focus shifts towards the full ‘interactome’ of organisms, PCR based approaches requiring multiple targets and primer sets are becoming less attractive. In addition, in many applications pools of specimens are more informative, such as to understand trophic interactions at the population level, mitigating some of this limitation.

No significant disadvantages of DDSS were evident compared to the current molecular methods in terms of PPV and taxonomic resolution. DDSS provides quantitative detection and identification of species taxa in degraded environmental DNA through a PCR-free method. This opens the possibility of quantifying food web interaction strengths without the complications of PCR-bias and errors that continue to plague metabarcoding, and will enable development of reliable, accurate empirical food webs that can be used to fuel new insights in community ecology.

## Data availability

The sequenced libraries (Fastq) were deposited at GenBank and their Sequence Read Archive (SRA) access codes are: SRR5342614 - SRR5342629 and SRR5350637-SRR5350656.

## Acknowledgements

We thank Lucas Machado de Souza and Bruna Lima for helping to collect ladybird beetles and aphids in the field; Professor Brígida Sousa from the Universidade Federal de Lavras (UFLA-Brazil) for supplying *Chrysoperla externa* eggs for our laboratory colony; and Embrapa and Graduate Program from the Department of Cell Biology for the PhD fellowships to RVT. We thank USDA-NIFA 2016-67030-24950 and USDA Regional Research Project NC-205 award for partial funding of this work.

## Author’s contributions

D.P.P., D.A.A., A.P.V.: conceptualization of the method; D.P.P., R.V.T., D.A.A.: design of study; R.V.T.: bioassays; R.V.T., D.P.P., R.C.T., D.A.A.: data analyses; D.P.P., R.C.T., A.P.V., D.A.A.: writing of the manuscript.

