## Supporting Information for "Quantitative prey species detection in predator guts across multiple trophic levels by DNA shotgun sequencing"

### Supplementary Information

Débora P. Paula[1,*], Renata V. Timbó [1,2], Roberto C. Togawa[1], Alfried P. Vogler [3,4] & David A. Andow [5]

*

[1]Embrapa Recursos Genéticos e Biotecnologia, Brasília, DF, Brazil

[2]Universidade de Brasília, Campus Universitário Darcy Ribeiro, Brasília, DF, Brazil

[3]Imperial College London, Silwood Park Campus, Ascot, UK

[4]Department of Life Sciences, Natural History Museum, London, UK

[5]Department of Entomology, University of Minnesota, St. Paul, USA

**Insect rearing**. *Harmonia axyridis* and *Hippodamia convergens* were reared in a climate-controlled chamber at 25±4°C, 60±20% RH and 16:8 h L:D. Field collected adults of the same species were maintained in three-liter plastic cages containing a daily supply of aphids from the species *Aphis gossypii*, *Myzus persicae* and *Brevicoryne brassicae* on their host plant leaves. Aphids were collected from organic farms in the Distrito Federal State (Brazil) twice a week and kept inside 500 ml plastic pots in a refrigerator (4°C) until used. Every other day the adult cages were cleaned (dry leaves without aphids were discarded) and inspected for the presence of egg masses. Egg masses were transferred to 50 ml plastic pots, and after larval eclosion and dispersal, they were separated individually in 50 ml plastic pots to prevent cannibalism, with a daily supply of aphids (one or more of the same species as for the adults rearing), and *Ephestia kuniella* eggs. Larval pots were also cleaned every other day until pupation. Newly emerged adults within 24 h were transferred to the adult cages. At least every three months the colonies received newly collected adults from the field. *Chrysoperla externa* rearing started with eggs obtained from the Entomology Laboratory at Universidade Federal de Lavras (UFLA), Lavras, Minas Gerais-Brazil, coordinated by Professor Dr. Brígida Souza. Chrysopid rearing was also performed in a climate-controlled chamber at 25±4°C, 60±20% RH and 12:12 h L:D. After eclosion, each larva was transferred individually to 50 ml plastic pots containing a daily supply of *E. kuehniella* and eventually aphids of the species *A. gossypii* and *M. persicae* until pupation. Newly emerged adults (ca, 20 individuals) within 24 h were transferred to PVC pipes (90 mm diameter, 30 cm height) lined with sulfite paper for oviposition. The cage pipes were closed at one end with a plastic screen and, at the other end, with organdy. Adult food consisted of a mixture of honey and beer yeast. Water was provided with a moistened cotton ball. Every other day the sulfite paper was replaced and eggs were transferred to 50 ml plastic pots. After larval eclosion, a daily supply of food was provided as described above.

**Prey detection threshold determination.** A Java program called BlastNToSnp (Lindenbaum 2015) was used to print indel/mismatch and gap relative positions in a BlastN stream. The BlastNToSnp was customized to add 100% matches in the output file. The modification of the original BlastNToSnp program, the in-house scripts to prepare datasets, and the *R* scripts are available in the GitHub repository: https://github.com/molecular-ecology/DDSS. The workflow continued using a series of customized *R* scripts: I) The output text file was modified to remove incompatible characters (e.g., commas) in the column names to run in *R* (script **correct text.*R***), and the BlastNToSnp output was analyzed to determine the absolute location of each potential mismatch between the read and its reference hit species (script **mismatch locations.*R***); II) The mismatch locations were examined to determine if the mismatch involved a degenerate base, and false mismatches were removed (script **match degenerate bases.*R***). This was necessary because BlastNToSNP considers matches to degenerate bases to be mismatches. Reads were filtered using 25 different thresholds, which comprised the combination of five overlap lengths (100, 125, 150, 175 and 200 bp) with five percent identity thresholds (96, 97, 98, 99 and 100%). These overlap lengths were used taking into consideration that the prey DNA is degraded in the gut content, therefore often with around 200 bp maximum (Zaidi et al. 1999; Agustí et al. 2003; Hoogendoorn & Heimpel 2003; Sheppard et al. 2004; Deagle et al. 2006). This was done by determining the highest percent identity between a read and the hit reference sequence for each hit match for a given threshold overlap length (script **percent identity.*R***). This was necessary because BlastN determines an overlap region and calculates the percent identity over this entire region. The BlastN match region can be longer than the overlap length of a threshold, and therefore, the percent identity for a shorter overlap length may differ from that for the entire BlastN match region (Fig. S1A). Then, because a single read typically matched more than one hit species, the hit species with the highest percent identity for a read among the set of hit matches for that read for a specified overlap length was identified (script **best matches.*R***). If the read had multiple reference hit species with the same highest percent identity, the names of all of these species were retained (these were called reads with multiple hit species). Reads with only one best reference hit match were called reads with single reference hit species. The number of reads for each hit match species and the number of distinct hit match species were counted for each threshold filter (overlap length and percent identity). Reads were filtered to retain only paired-end reads with identical hit match species for each specified overlap length and percent identity, and the best thresholds were identified. Potentially acceptable thresholds were those that met at least one of three criteria: a) eliminated at least 1/3 of false positive reads while retaining at least ¾ of the true positive prey reads; b) eliminated at least 1/3 of false positive reads while retaining at least 1/2 of the true positive predator reads; or c) eliminated half of the false positive species while retaining 80% of the true positive prey or eliminated 2/3 of the false positive species while retaining ¾ of the true positive prey. The best thresholds were ones that met a higher number of these criteria. The workflow continued by mapping the remaining read sequences associated with the best thresholds to the mitogenome of the identified hit species using Geneious Prime 2 2020 mapper with the parameters: custom sensitivity, do not trim, minimum mapping quality 30, allow gaps, maximum per read 20%, maximum gap size 500, minimum overlap length 100 to 175 bp, minimum overlap identity 99 to 100%, word length 10, index word length 10, ignore words repeated more than 100 times, maximum mismatches per read 50%, maximum ambiguity 16. After mapping, only paired-end reads were retained. Finally, the number of reads for each species in the control libraries (predator only, no prey consumed) was used as a blank to subtract from all of the experimental libraries.

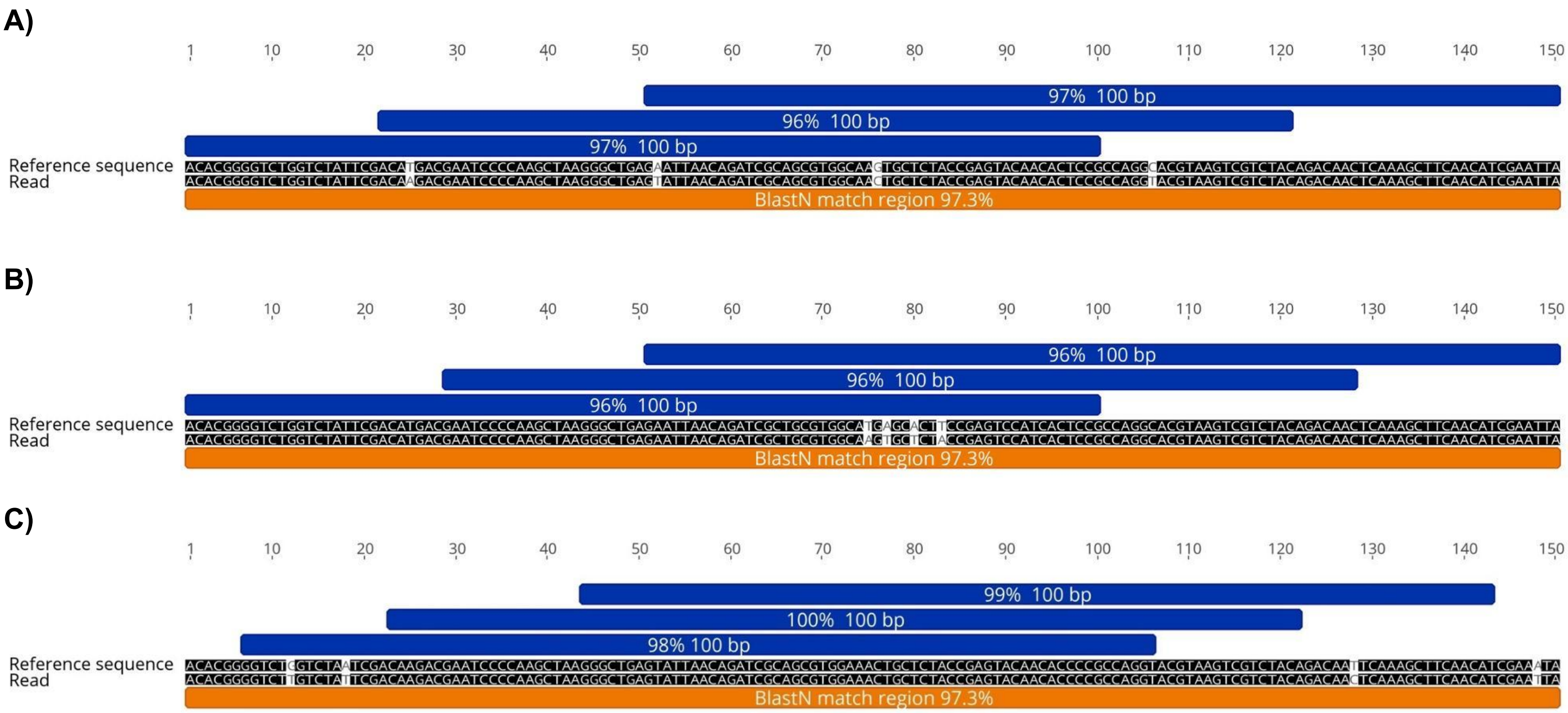


**Fig. S1A** Illustration of the importance of the mismatch positions in a BlastN match region (150 bp) to filter for false positive detection. A) to C) represent an alignment between a reference sequence and a read containing four mismatches, both resulting in 97.3% identity for the entire match. In A) all possible 100 bp overlap lengths contain 3-4 mismatches and have 96% or 97% identity. This read would satisfy the threshold 100bp_97%. In B) all possible 100 bp overlap lengths contain 4 mismatches, and this read would not satisfy the threshold 100bp_97%. In C) there are regions in the BlastN match region with 100 bp overlap that contain 0, 1 or 2 mismatches with identities of 100%, 99% and 98%, respectively. Thus, this read would satisfy filtering thresholds of all the thresholds (100bp_98%, 100bp_99% and 100bp_100%).

**Regressions:**

**1) Effect of prey number.** Individual regressions were done for each prey number ingested as described above. The estimated parameter values (ln($n_0$), *d*, and $D_{max}$) for the three regressions were compared by ANOVA using the standard errors of the parameters to estimate the error mean square. As prey reads followed a first-order decay process, this implied that the ln of ingested prey number should be directly proportional to the ln of detected prey reads, so a multiple regression of time and ln ingested prey number on ln prey reads was conducted. We tested the prediction of direct proportionality by testing if the regression parameter was different from 1. Finally, $D_{max}$ was also predicted to be a function of the ln of ingested prey number, so we fit this model to the estimated $D_{max}$.

**2) Decay of *M. persicae* in the secondary predator, *Ha. axyridis*.** Three *M. persicae* were ingested by a *C. externa* larva, and 0, 3 or 6 h later, the *C. externa* larva was ingested by *Ha. axyridis*, and the number of *M. persicae* reads detected 0, 3 or 6 h later were estimated. Thus, secondary decay in *Ha. axyridis* was estimated controlling for the time after ingestion by *C. externa*. In addition, *Ha. axyridis* directly ingested *M. persicae* to compare direct and indirect detection and decay.

**3) Decay of *M. persicae* reads in the primary predator, *C. externa* larva.** This used the same data as in 2), except that secondary decay in *C. externa* was estimated controlling for the time after ingestion by *Ha. axyridis*. We also determined if decay of *M. persicae* in *C. externa* was similar to decay in *Ha. axyridis*. This was done by comparing primary decay in *C. externa* (direct predation) with primary decay in *Ha. axyridis* (direct predation), and by comparing primary decay in *C. externa* (direct predation) with secondary decay in *Ha. axyridis* (indirect predation, from 2 above).

**4) Decay of *C. externa* reads in the predator, *Ha. axyridis*.** One *C. externa* larva was ingested by *Ha. axyridis* and the number of *C. externa* reads in *Ha. axyridis* was estimated 0, 3 and 6 h after ingestion. The *C. externa* larvae were either unfed, or had fed on three *M. persicae* 0, 3 or 6 h before being ingested by *Ha. axyridis*. We tested if the decay rate depended on the feeding history of *C. externa*. We also tested if the number of *M. persicae* and *C. externa* reads were correlated among the libraries where both were detected.

**Prey detectability**

All of the statistical analyses had similar results for all of the best six thresholds. The results for the four sets of regression mentioned in the Statistical Methods section are presented as follows.

*1) Effect of prey number.* The decay of *M. persicae* reads in *Hi. convergens* after ingestion of 1, 3 or 6 *M. persicae* apterae followed a first order decay process and the results were similar for all six thresholds (Fig. S1). In all cases there was no difference in the decay rate, but $\ln(n_0)$ was higher and $D_{max}$ was longer when more prey were consumed (Tables S1-S2). The number of prey reads detected was directly related to the number of prey ingested (parameter estimate for $\ln(Mp)$ was not different from 1 (Table S3).

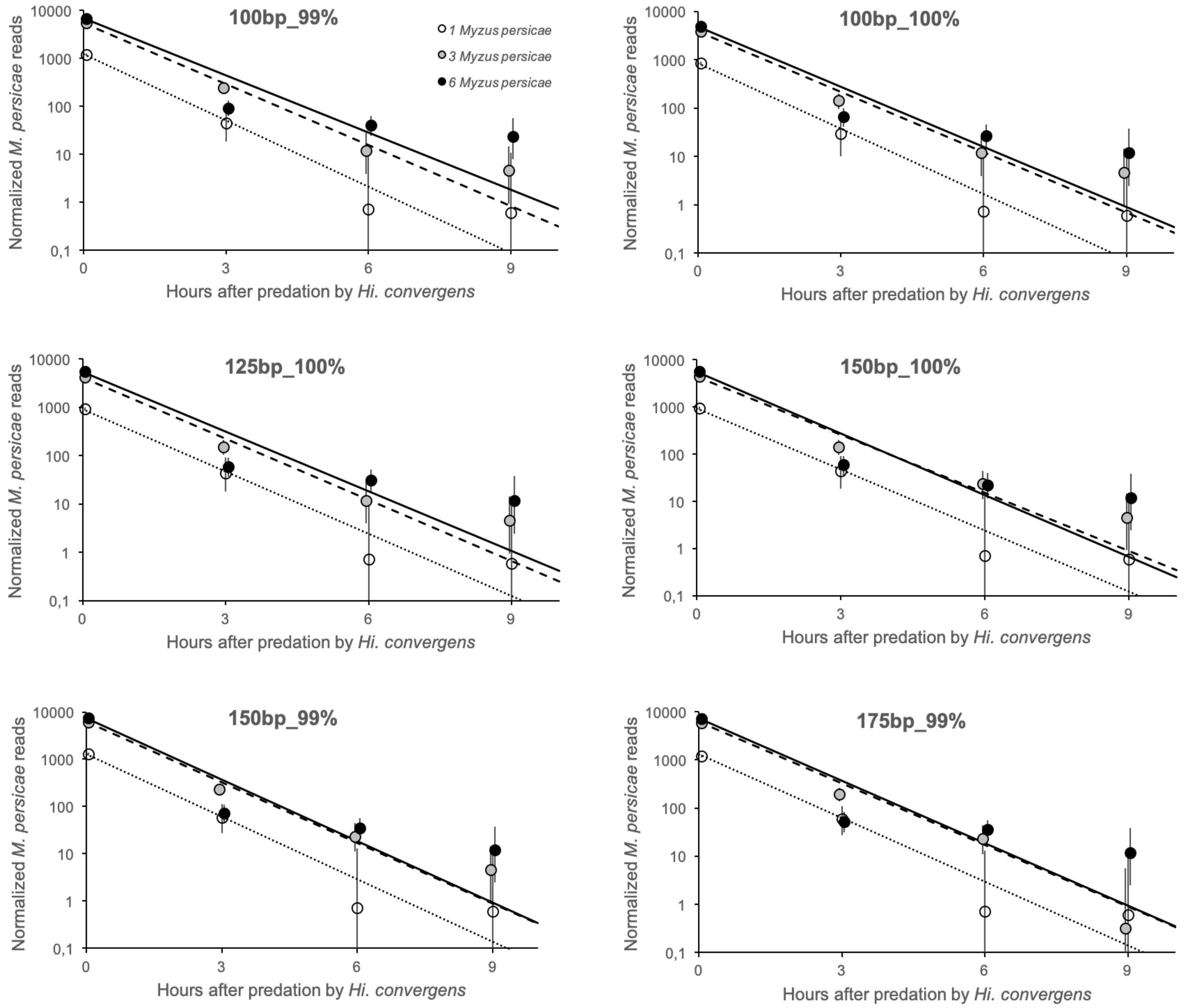


**Figure S1.** Decay of *Myzus persicae* reads in *Hippodamia convergens* after ingestion of 1, 3 or 6 *M. persicae* apterae for the six best thresholds (overlap length_%identity).

**Table S1.** Estimated parameter values using the six best thresholds (overlap length_%identity) for the detection of different numbers of *Myzus persicae* prey ingested by *Hippodamia convergens* larvae (means with 95% CIs in brackets). $n_0$: initial prey reads, $d$: decay rate, $D_{max}$: maximum detectability period.

| Threshold | Prey | ln($n_0$) | $d$ ($h^{-1}$) | $D_{max}$ (h) |
|---|---|---|---|---|
| 100_100 | 1 *M. persicae* | 6.73 (0.03) | -1.04 (0.10) | 7.54 (0.60) |
| | 3 *M. persicae* | 8.23 (0.10) | -0.96 (0.11) | 10.27 (0.40) |
| | 6 *M. persicae* | 8.48 (0.14) | -0.95 (0.20) | 11.92 (0.54) |
| 100_99 | 1 *M. persicae* | 7.09 (0.03) | -1.06 (0.08) | 7.57 (0.57) |
| | 3 *M. persicae* | 8.59 (0.07) | -0.98 (0.07) | 10.15 (0.37) |
| | 6 *M. persicae* | 8.81 (0.15) | -0.91 (0.20) | 13.08 (0.47) |
| 125_100 | 1 *M. persicae* | 6.83 (0.03) | -0.99 (0.07) | 7.60 (0.60) |
| | 3 *M. persicae* | 8.33 (0.10) | -0.97 (0.11) | 10.23 (0.39) |
| | 6 *M. persicae* | 8.59 (0.14) | -0.95 (0.20) | 11.96 (0.54) |
| 150_100 | 1 *M. persicae* | 6.84 (0.03) | -0.99 (0.07) | 7.60 (0.60) |
| | 3 *M. persicae* | 8.39 (0.12) | -0.94 (0.11) | 10.60 (0.40) |
| | 6 *M. persicae* | 8.62 (0.13) | -1.00 (0.21) | 11.67 (0.52) |
| 150_99 | 1 *M. persicae* | 7.16 (0.02) | -1.02 (0.07) | 7.61 (0.57) |
| | 3 *M. persicae* | 8.72 (0.08) | -0.98 (0.08) | 10.48 (0.37) |
| | 6 *M. persicae* | 8.91 (0.13) | -1.00 (0.21) | 11.82 (0.51) |
| 175_99 | 1 *M. persicae* | 7.11 (0.02) | -1.00 (0.07) | 7.62 (0.57) |
| | 3 *M. persicae* | 8.67 (0.07) | -1.04 (0.08) | 8.53 (0.60) |
| | 6 *M. persicae* | 8.86 (0.13) | -0.97 (0.21) | 11.85 (0.52) |

**Table S2.** ANOVA for regression parameters for the six best thresholds (overlap length_%identity). *Mp* is the number of *Myzus persicae* ingested.

| **Threshold** | | **ln($n_0$)** | | | | ***d* ($h^{-1}$)** | | | | **$D_{max}$ (h)** | | | |
|---|---|---|---|---|---|---|---|---|---|---|---|---|---|
| | **Factor** | ***df*** | ***MS*** | ***F*** | ***p*** | ***df*** | ***MS*** | ***F*** | ***p*** | ***df*** | ***MS*** | ***F*** | ***p*** |
| 100_100 | Mp | 2 | 0.4448 | 43.53 | 2.359E-5 | 2 | 0.00117 | 0.058 | 0.9436 | 2 | 2.455 | 9.02 | 0.0071 |
| | Error | 9 | 0.0102 | | | 9 | 0.02003 | | | 9 | 0.272 | | |
| 100_99 | Mp | 2 | 0.4386 | 48.07 | 1.571E-5 | 2 | 0.00278 | 0.158 | 0.8561 | 2 | 3.800 | 16.67 | 0.0009 |
| | Error | 9 | 0.0091 | | | 9 | 0.01756 | | | 9 | 0.228 | | |
| 125_100 | Mp | 2 | 0.4531 | 45.98 | 1.885E-5 | 2 | 0.00021 | 0.011 | 0.9892 | 2 | 2.402 | 8.98 | 0.0072 |
| | Error | 9 | 0.0099 | | | 9 | 0.01901 | | | 9 | 0.267 | | |
| 150_100 | Mp | 2 | 0.4723 | 45.66 | 1.940E-5 | 2 | 0.00047 | 0.022 | 0.9782 | 2 | 2.232 | 8.50 | 0.0084 |
| | Error | 9 | 0.0103 | | | 9 | 0.02137 | | | 9 | 0.262 | | |
| 150_99 | Mp | 2 | 0.4610 | 59.51 | 6.475E-6 | 2 | 0.00016 | 0.009 | 0.9913 | 2 | 2.309 | 9.61 | 0.0058 |
| | Error | 9 | 0.0077 | | | 9 | 0.01806 | | | 9 | 0.240 | | |
| 175_99 | Mp | 2 | 0.4623 | 62.79 | 5.171E-6 | 2 | 0.00055 | 0.029 | 0.9711 | 2 | 2.475 | 7.77 | 0.0109 |
| | Error | 9 | 0.0074 | | | 9 | 0.01864 | | | 9 | 0.318 | | |

**Table S3.** Parameter estimates for the six best thresholds (overlap length_%identity) for multiple regression model, $\ln(n(T, Mp)) = \ln(n_0) + m_1 \ln(Mp) + m_2 T$, where $\ln(n(T, Mp))$ is the ln-transform of the number of prey reads (for time $T$, and $Mp$ prey ingested), $Mp$ is the number of *Myzus persicae* ingested, $T$ is the time after consumption, $m_1$ and $m_2$ are the regression slopes, and $n_0$ is the initial number of prey reads (time = 0 and for 1 prey).

| | | **test: ln(*Mp*) = 0** | | | | **test: ln(*Mp*) = 1** | | |
|---|---|---|---|---|---|---|---|---|
| **Threshold** | **Parameter** | **Estimate** | **SEM** | ***t*-value** | ***p*** | **Estimate** | ***t*-value** | ***p*** |
| 100_100 | ln($n_0$) | 6.861641 | 0.16304 | 42.08553 | 1.20E-11 | | | |
| | time | -0.93774 | 0.11194 | -8.37678 | 1.53E-05 | | | |
| | ln(Mp) | 0.941786 | 0.11187 | 8.418584 | 1.47E-05 | 0.058214 | 0.520371 | 0.615363 |
| 100_99 | ln($n_0$) | 7.218495 | 0.16526 | 43.67756 | 8.63E-12 | | | |
| | time | -0.91891 | 0.11290 | -8.13849 | 1.93E-05 | | | |
| | ln(Mp) | 0.93305 | 0.11370 | 8.206094 | 1.81E-05 | 0.06695 | 0.588824 | 0.57046 |
| 125_100 | ln($n_0$) | 6.950533 | 0.15892 | 43.73616 | 8.53E-12 | | | |
| | time | -0.94009 | 0.11203 | -8.39121 | 1.51E-05 | | | |
| | ln(Mp) | 0.955849 | 0.10885 | 8.780712 | 1.04E-05 | 0.044151 | 0.40558 | 0.694524 |
| 150_100 | ln($n_0$) | 6.979959 | 0.17185 | 40.6155 | 1.66E-11 | | | |
| | time | -0.94668 | 0.12242 | -7.73301 | 2.90E-05 | | | |
| | ln(Mp) | 0.960197 | 0.11762 | 8.163187 | 1.88E-05 | 0.039803 | 0.338389 | 0.74283 |
| 150_99 | ln($n_0$) | 7.309938 | 0.16901 | 43.25127 | 9.43E-12 | | | |
| | time | -0.96041 | 0.12432 | -7.72498 | 2.92E-05 | | | |
| | ln(Mp) | 0.94175 | 0.11629 | 8.098281 | 2.01E-05 | 0.05825 | 0.500899 | 0.628463 |
| 175_99 | ln($n_0$) | 7.254733 | 0.16689 | 43.46904 | 9.01E-12 | | | |
| | time | -0.97372 | 0.12986 | -7.49814 | 3.70E-05 | | | |
| | ln(Mp) | 0.945472 | 0.11486 | 8.231082 | 1.76E-05 | 0.054528 | 0.47471 | 0.646301 |

*2) Decay of* M. persicae *in the secondary predator,* Ha. axyridis. The decay of *M. persicae* reads after indirect predation by *Ha. axyridis* followed a first order decay process and the results were similar for all six thresholds (Fig. S2). There were no significant differences in the decay rates for direct versus indirect predation of *M. persicae* by *Ha. axyridis*. However, more *M. persicae* reads were detected after direct ingestion than for indirect ingestion (Tables S4 and S5).

**Table S4.** Estimated parameter values using the six best thresholds (overlap length_%identity) for the detection of *Chrysoperla externa* (Ce) and *Myzus persicae* (Mp) consumed directly or indirectly (via predation on Ce) by *Harmonia axyridis* (Ha) larvae (means with 95% CIs in brackets). D= direct predation; I= indirect predation.

| Threshold | Predator | Prey | Feeding history | Time (h) after direct Mp consumption by Ha | Time (h) after direct Mp consumption by Ce | Time (h) after direct Ce consumption by Ha | ln($n_0$) | *d* ($h^{-1}$) | $D_{max}$ (h) |
|---|---|---|---|---|---|---|---|---|---|
| 100_100 | **Predation on Ce** | | | | | | | | |
| | Ha (D) | 1Ce | None | -- | -- | 0-6 | 7.59 (0.08) | -0.43 (0.04) | 18.98 (0.75) |
| | Ha (D) | 1Ce | 3Mp | -- | 0 | 0-6 | 5.26 (0.25) | -0.43 (0.15) | 6.72 (1.14) |
| | Ha (D) | 1Ce | 3Mp | -- | 3 | 0-6 | 5.42 (0.25) | -0.25 (0.10) | 25.29 (3.25) |
| | Ha (D) | 1Ce | 3Mp | -- | 6 | 0-6 | 4.11 (0.01) | -0.97 (0.01) | 4.72 (0.76) |
| | **Predation on Mp** | | | | | | | | |
| | Ha (D) | 3Mp | None | 0-6 | -- | -- | 7.48 (0.35) | -0.29 (0.16) | 13.49 (0.73) |
| | Ha (I) | 3Mp in Ce | None | -- | 0 | 0-6 | 4.31 (0.23) | -0.13 (0.07) | -- |
| | Ha (I) | 3Mp in Ce | None | -- | 3 | 0-6 | 4.18 (0.10) | -0.62 (0.07) | 6.89 (1.83) |
| | Ha (I) | 3Mp in Ce | None | -- | 6 | 0-6 | -0.84 (0.24) | -0.15 (0.06) | -- |
| | Ce (D) | 3Mp in Ce | None | -- | 0-6 | 0 | 4.22 (0.15) | -0.65 (0.09) | 7.08 (1.57) |
| | Ce (D) | 3Mp in Ce | None | -- | 0-6 | 3 | 3.34 (0.18) | -0.89 (0.18) | 5.33 (0.91) |
| | Ce (D) | 3Mp in Ce | None | -- | 0-6 | 6 | 4.54 (1.56) | -0.40 (0.62) | 5.91 (0.99) |
| 100_99 | **Predation on Ce** | | | | | | | | |
| | Ha (D) | 1Ce | None | -- | -- | 0-6 | 8.92 (0.10) | -0.49 (0.05) | 19.81 (0.41) |
| | Ha (D) | 1Ce | 3Mp | -- | 0 | 0-6 | 6.81 (0.16) | -0.48 (0.08) | 11.92 (0.73) |
| | Ha (D) | 1Ce | 3Mp | -- | 3 | 0-6 | 6.80 (0.17) | -0.26 (0.07) | 28.66 (1.64) |
| | Ha (D) | 1Ce | 3Mp | -- | 6 | 0-6 | 4.65 (0.29) | -0.71 (0.17) | 7.4 0(0.73) |
| | **Predation on Mp** | | | | | | | | |
| | Ha (D) | 3Mp | None | 0-6 | -- | -- | 7.79 (0.31) | -0.33 (0.14) | 14.27 (0.65) |
| | Ha (I) | 3Mp in Ce | None | -- | 0 | 0-6 | 4.85 (0.10) | -0.20 (0.03) | 26.63 (53.77) |
| | Ha (I) | 3Mp in Ce | None | -- | 3 | 0-6 | 4.50 (0.13) | -0.60 (0.09) | 6.83 (2.83) |
| | Ha (I) | 3Mp in Ce | None | -- | 6 | 0-6 | -0.84 (0.23) | -0.15 (0.06) | 9.09 (1744) |

| | | | | | | | | | |
|---|---|---|---|---|---|---|---|---|---|
| | Ce (D) | 3Mp in Ce | None | -- | 0-6 | 0 | 4.82 (0.14) | -0.70 (0.09) | 6.83 (1.22) |
| | Ce (D) | 3Mp in Ce | None | -- | 0-6 | 3 | 3.56 (0.18) | -0.94 (0.19) | 5.44 (0.88) |
| | Ce (D) | 3Mp in Ce | None | -- | 0-6 | 6 | 4.79 (1.61) | -0.37 (0.64) | 5.96 (0.95) |
| 125_100 | **Predation on Ce** | | | | | | | | |
| | Ha (D) | 1Ce | None | -- | -- | 0-6 | 7.65 (0.08) | -0.44 (0.04) | 18.67 (0.72) |
| | Ha (D) | 1Ce | 3Mp | -- | 0 | 0-6 | 5.48 (0.13) | -0.47 (0.06) | 10.56 (1.20) |
| | Ha (D) | 1Ce | 3Mp | -- | 3 | 0-6 | 5.49 (0.24) | -0.24 (0.09) | 26.73 (3.38) |
| | Ha (D) | 1Ce | 3Mp | -- | 6 | 0-6 | 4.14 (0.20) | -0.66 (0.13) | 5.15 (0.83) |
| | **Predation on Mp** | | | | | | | | |
| | Ha (D) | 3Mp | None | 0-6 | -- | -- | 7.60 (0.33) | -0.31 (0.15) | 13.22 (0.69) |
| | Ha (I) | 3Mp in Ce | None | -- | 0 | 0-6 | 4.19 (0.03) | -0.07 (0.01) | -- |
| | Ha (I) | 3Mp in Ce | None | -- | 3 | 0-6 | 4.18 (0.10) | -0.62 (0.07) | 6.90 (1.87) |
| | Ha (I) | 3Mp in Ce | None | -- | 6 | 0-6 | -0.84 (0.23) | -0.15 (0.06) | -- |
| | Ce (D) | 3Mp in Ce | None | -- | 0-6 | 0 | 4.22 (0.16) | -0.64 (0.10) | 7.09 (1.55) |
| | Ce (D) | 3Mp in Ce | None | -- | 0-6 | 3 | 3.74 (0.18) | -0.97 (0.20) | 5.36 (0.96) |
| | Ce (D) | 3Mp in Ce | None | -- | 0-6 | 6 | 4.43 (1.66) | -0.37 (0.64) | 5.97 (1.05) |
| 150_100 | **Predation on Ce** | | | | | | | | |
| | Ha (D) | 1Ce | None | -- | -- | 0-6 | 7.66 (0.10) | -0.43 (0.05) | 19.35 (0.73) |
| | Ha (D) | 1Ce | 3Mp | -- | 0 | 0-6 | 5.43 (0.14) | -0.46 (0.07) | 10.64 (1.23) |
| | Ha (D) | 1Ce | 3Mp | -- | 3 | 0-6 | 5.58 (0.15) | -0.27 (0.06) | 23.51 (2.56) |
| | Ha (D) | 1Ce | 3Mp | -- | 6 | 0-6 | 4.14 (0.20) | -0.66 (0.13) | 5.15 (0.83) |
| | **Predation on Mp** | | | | | | | | |
| | Ha (D) | 3Mp | None | 0-6 | -- | -- | 7.58 (0.34) | -0.30 (0.16) | 13.28 (0.70) |
| | Ha (I) | 3Mp in Ce | None | -- | 0 | 0-6 | 4.30 (0.20) | -0.07 (0.05) | -- |
| | Ha (I) | 3Mp in Ce | None | -- | 3 | 0-6 | 4.05 (0.12) | -0.58 (0.08) | 7.03 (5.18) |
| | Ha (I) | 3Mp in Ce | None | -- | 6 | 0-6 | -0.84 (0.24) | -0.15 (0.06) | -- (--) |
| | Ce (D) | 3Mp in Ce | None | -- | 0-6 | 0 | 4.22 (0.16) | -0.64 (0.10) | 7.03 (1.54) |
| | Ce (D) | 3Mp in Ce | None | -- | 0-6 | 3 | 3.74 (0.17) | -0.97 (0.20) | 5.31 (0.88) |
| | Ce (D) | 3Mp in Ce | None | -- | 0-6 | 6 | 4.61 (1.40) | -0.47 (0.57) | 5.82 (0.95) |
| 150_99 | **Predation on Ce** | | | | | | | | |
| | Ha (D) | 1Ce | None | -- | -- | 0-6 | 8.89 (0.09) | -0.49 (0.05) | 19.70 (0.41) |
| | Ha (D) | 1Ce | 3Mp | -- | 0 | 0-6 | 6.80 (0.10) | -0.49 (0.05) | 12.83 (0.76) |

| | | | | | | | | | |
|---|---|---|---|---|---|---|---|---|---|
| | Ha (D) | 1Ce | 3Mp | -- | 3 | 0-6 | 6.79 (0.09) | -0.29 (0.04) | 25.01 (1.37) |
| | Ha (D) | 1Ce | 3Mp | -- | 6 | 0-6 | 4.60 (0.19) | -0.67 (0.10) | 7.68 (0.77) |
| | **Predation on Mp** | | | | | | | | |
| | Ha (D) | 3Mp | None | 0-6 | -- | -- | 7.80 (0.32) | -0.32 (0.15) | 14.27 (0.65) |
| | Ha (I) | 3Mp in Ce | None | -- | 0 | 0-6 | 4.67 (0.14) | -0.11 (0.04) | -- |
| | Ha (I) | 3Mp in Ce | None | -- | 3 | 0-6 | 4.18 (0.10) | -0.62 (0.07) | 6.89 (1.92) |
| | Ha (I) | 3Mp in Ce | None | -- | 6 | 0-6 | -0.85 (0.24) | -0.15 (0.06) | -- |
| | Ce (D) | 3Mp in Ce | None | -- | 0-6 | 0 | 4.62 (0.09) | -0.77 (0.06) | 6.81 (1.29) |
| | Ce (D) | 3Mp in Ce | None | -- | 0-6 | 3 | 3.90 (0.17) | -1.00 (0.21) | 5.28 (0.83) |
| | Ce (D) | 3Mp in Ce | None | -- | 0-6 | 6 | 4.80 (1.36) | -0.48 (0.56) | 5.81 (0.90) |
| 175_99 | **Predation on Ce** | | | | | | | | |
| | Ha (D) | 1Ce | None | -- | -- | 0-6 | 8.79 (0.07) | -0.49 (0.04) | 19.13 (0.43) |
| | Ha (D) | 1Ce | 3Mp | -- | 0 | 0-6 | 6.67 (0.08) | -0.45 (0.04) | 14.05 (0.84) |
| | Ha (D) | 1Ce | 3Mp | -- | 3 | 0-6 | 6.74 (0.15) | -0.26 (0.06) | 28.34 (1.68) |
| | Ha (D) | 1Ce | 3Mp | -- | 6 | 0-6 | 4.61 (0.10) | -0.64 (0.05) | 7.85 (0.78) |
| | **Predation on Mp** | | | | | | | | |
| | Ha (D) | 3Mp | None | 0-6 | -- | -- | 7.73 (0.32) | -0.32 (0.15) | 12.97 (0.65) |
| | Ha (I) | 3Mp in Ce | None | -- | 0 | 0-6 | 4.42 (0.39) | -0.08 (0.11) | -- (--) |
| | Ha (I) | 3Mp in Ce | None | -- | 3 | 0-6 | 4.16 (0.03) | -0.80 (0.03) | 6.54 (5.79) |
| | Ha (I) | 3Mp in Ce | None | -- | 6 | 0-6 | -0.83 (0.23) | -0.15 (0.06) | 3.07 (237.0) |
| | Ce (D) | 3Mp in Ce | None | -- | 0-6 | 0 | 4.20 (0.01) | -0.83 (0.01) | 7.09 (1.56) |
| | Ce (D) | 3Mp in Ce | None | -- | 0-6 | 3 | 3.74 (0.17) | -0.97 (0.20) | 5.01 (0.78) |
| | Ce (D) | 3Mp in Ce | None | -- | 0-6 | 6 | 4.80 (1.36) | -0.48 (0.56) | 5.81 (0.90) |

**Table S5.** ANOVA comparing *Myzus persicae* read detection in *Harmonia axyridis* after direct versus indirect predation for the six best thresholds (overlap length_%identity).

| **Threshold** | | **100_100** | | | **100_99** | | | **125_100** | | |
|---|---|---|---|---|---|---|---|---|---|---|
| | ***df*** | ***MS*** | ***F*** | ***p*** | ***MS*** | ***F*** | ***p*** | ***MS*** | ***F*** | ***p*** |
| Direct versus indirect | 1 | 6.929 | 100.53 | 0.00017 | 6.633 | 136.56 | 0.00008 | 6.764 | 105.94 | 0.00015 |
| Time | 2 | 2.679 | 38.87 | 0.00090 | 2.885 | 59.39 | 0.00033 | 2.233 | 34.98 | 0.00115 |
| Interaction | 2 | 0.965 | 13.999 | 0.00894 | 0.687 | 14.153 | 0.00873 | 1.452 | 22.747 | 0.00309 |
| Error | 5 | 0.069 | | | 0.049 | | | 0.064 | | |

| **Threshold** | | **150_100** | | | **150_99** | | | **175_99** | | |
|---|---|---|---|---|---|---|---|---|---|---|
| | ***df*** | ***MS*** | ***F*** | ***p*** | ***MS*** | ***F*** | ***p*** | ***MS*** | ***F*** | ***p*** |
| Direct versus indirect | 1 | 6.160 | 102.18 | 0.00016 | 6.227 | 139.62 | 0.00008 | 6.227 | 106.93 | 0.00015 |
| Time | 2 | 2.334 | 38.71 | 0.00091 | 2.335 | 52.36 | 0.00044 | 2.616 | 44.93 | 0.00064 |
| Interaction | 2 | 1.352 | 22.424 | 0.00319 | 1.019 | 22.845 | 0.00306 | 1.433 | 24.608 | 0.00258 |
| Error | 5 | 0.060 | | | 0.045 | | | 0.058 | | |

**Table S6.** ANOVA of decay rate of *Myzus persicae* in *Harmonia axyridis* after direct versus indirect predation for the six best thresholds (overlap length_%identity).

| **Threshold** | | ***df*** | ***SS*** | ***MS*** | ***F*** | ***p*** |
|---|---|---|---|---|---|---|
| 100_100 | Treatment | 3 | 0.1543 | 0.0514 | 3.95 | 0.1446 |
| | Direct vs indirect | 1 | 0.0125 | 0.0125 | 0.96 | 0.4000 |
| | All indirect | 2 | 0.1543 | 0.0771 | 5.92 | 0.0910 |
| | Error | 3 | 0.0391 | 0.0130 | | |
| 100_99 | Treatment | 3 | 0.1205 | 0.0402 | 3.58 | 0.1614 |
| | Direct vs indirect | 1 | 0.0080 | 0.0080 | 0.72 | 0.4593 |
| | All indirect | 2 | 0.1204 | 0.0602 | 5.36 | 0.1021 |
| | Error | 3 | 0.0337 | 0.0112 | | |
| 125_100 | Treatment | 3 | 0.1772 | 0.0591 | 5.52 | 0.0971 |
| | Direct vs indirect | 1 | 0.0283 | 0.0283 | 2.65 | 0.2023 |
| | All indirect | 2 | 0.1765 | 0.0882 | 8.25 | 0.0604 |
| | Error | 3 | 0.0321 | 0.0107 | | |
| 150_100 | Treatment | 3 | 0.1511 | 0.0504 | 3.98 | 0.1432 |
| | Direct vs indirect | 1 | 0.0255 | 0.0255 | 2.02 | 0.2506 |
| | All indirect | 2 | 0.1503 | 0.0752 | 5.94 | 0.0906 |
| | Error | 3 | 0.0380 | 0.0127 | | |
| 150_99 | Treatment | 3 | 0.1620 | 0.0540 | 4.99 | 0.1097 |
| | Direct vs indirect | 1 | 0.0205 | 0.0205 | 1.89 | 0.2627 |
| | All indirect | 2 | 0.1617 | 0.0808 | 7.47 | 0.0683 |
| | Error | 3 | 0.0324 | 0.0108 | | |
| 175_99 | Treatment | 3 | 0.3187 | 0.1062 | 8.23 | 0.0586 |
| | Direct vs indirect | 1 | 0.0297 | 0.0297 | 2.30 | 0.2266 |
| | All indirect | 2 | 0.3183 | 0.1591 | 12.32 | 0.0357 |
| | Error | 3 | 0.0387 | 0.0129 | | |

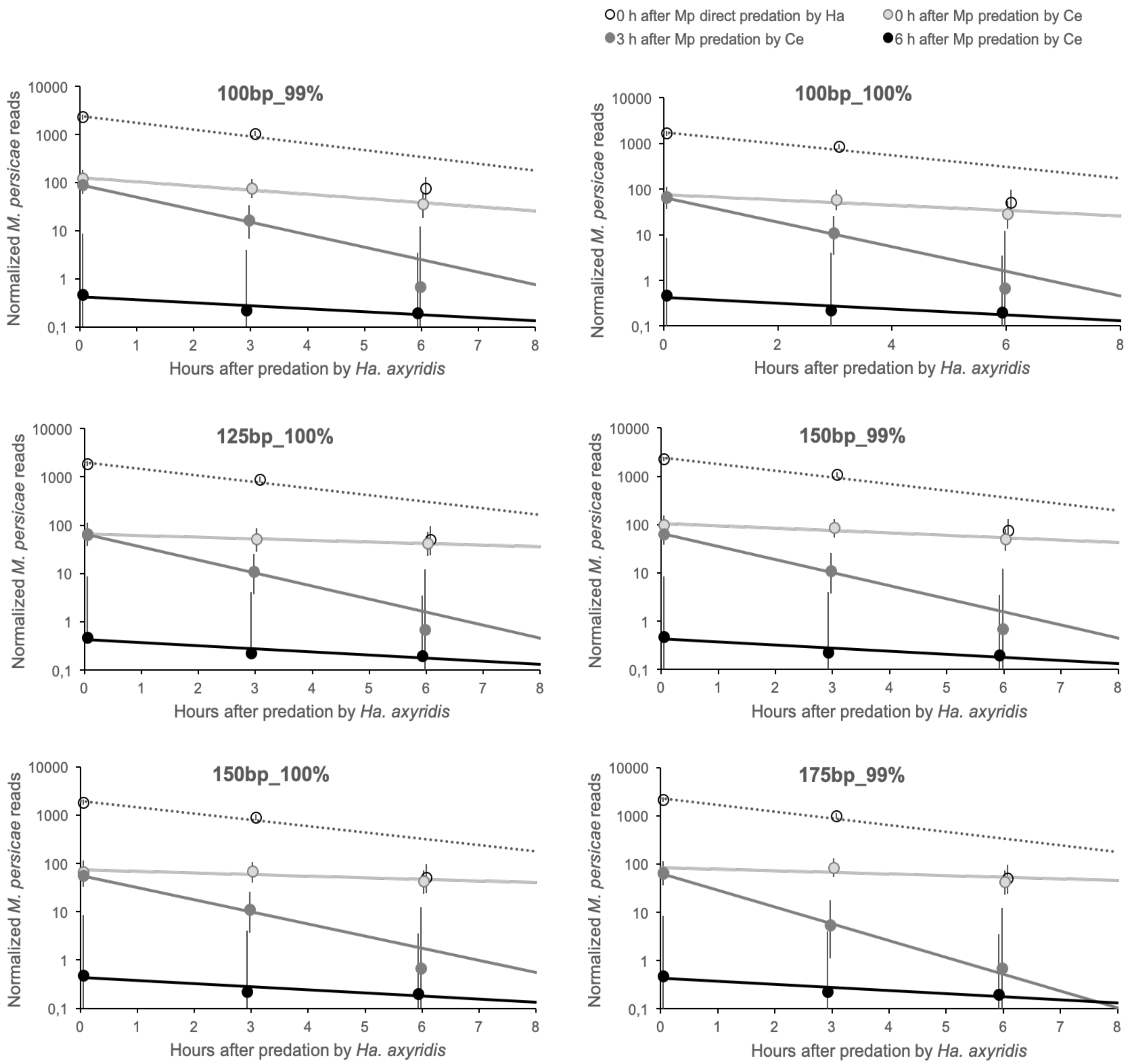


**Figure S2.** Indirect predation. Decay of *Myzus persicae* reads in the secondary predator, *Harmonia axyridis*, after ingestion of *Chrysoperla externa* that had ingested three *M. persicae* apterae for the six best thresholds (overlap length_%identity).

*3) Decay of* M. persicae *reads in the primary predator,* C. externa *larva.* The decay of *M. persicae* reads after ingestion by the primary predator, *C. externa*, followed a first order decay process and the results were similar for all six thresholds (Fig. S3). There were no significant differences in the decay rates of *M. persicae* after direct ingestion by *Ha. axyridis* versus after ingestion by *C. externa*, which was immediately ingested by *Ha. axyridis* (Table S6, direct vs indirect). In addition, there were no significant differences in decay rates depending on the time after *C. externa* was ingested by *Ha. axyridis* (Table S6, all indirect). Finally, we compared the secondary decay of *M. persicae* in *Ha. axyridis* with the decay of *M. persicae* in *C. externa*, which was itself ingested by *Ha. axyridis*, and found that the decay rates were similar (Table S7), i.e., the decay rate of *M. persicae* in *Ha. axyridis* was not significantly different from its decay rate in *C. externa*.

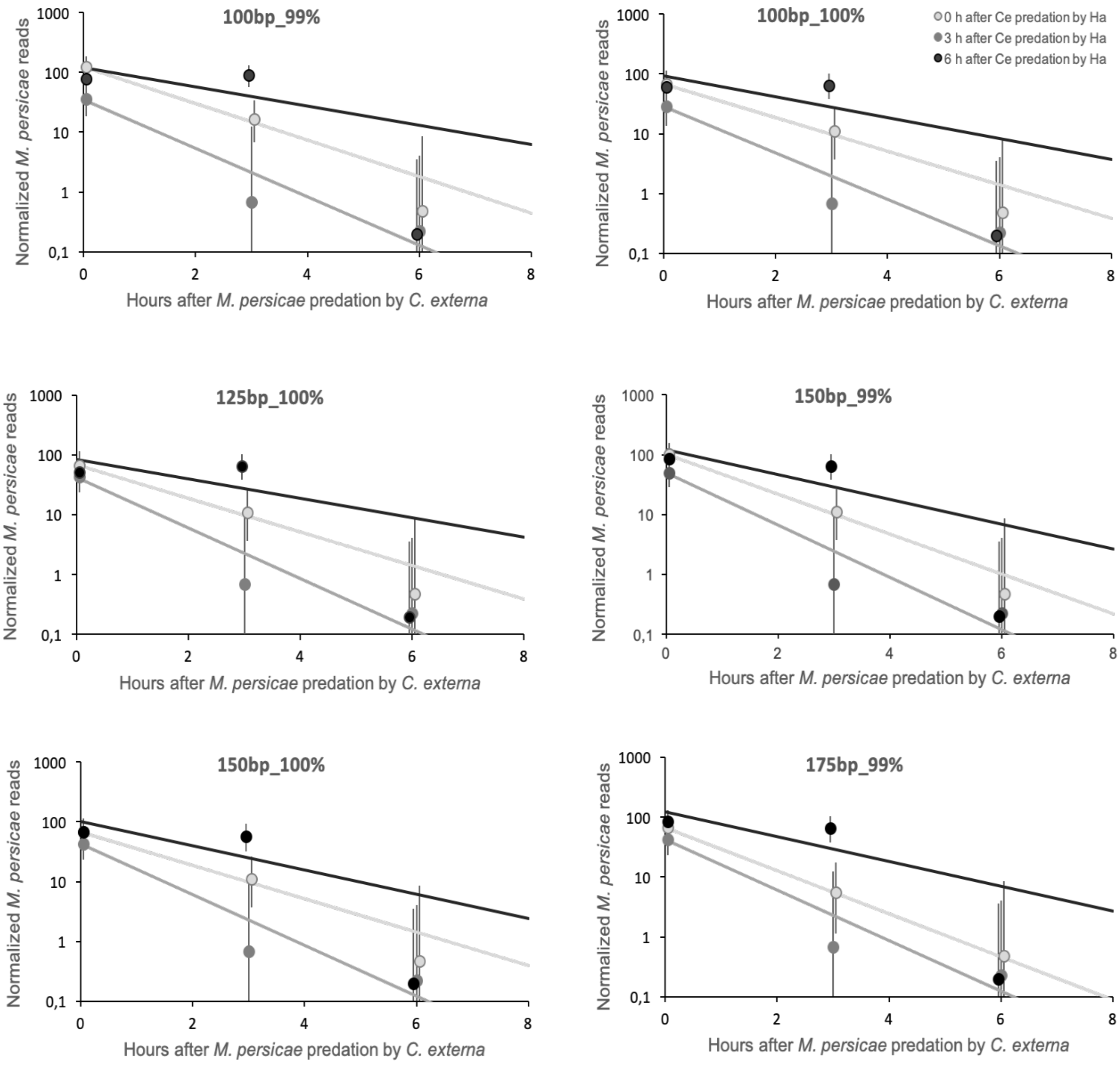


**Figure S3.** Direct predation. Decay of *Myzus persicae* reads in the primary predator, *Chrysoperla externa*, prior to ingestion of *C. externa* by *Harmonia axyridis* for the six best thresholds (overlap length_%identity).

**Table S7.** ANOVA of decay rate of *Myzus persicae* after indirect predation by *Harmonia axyridis* (Ha) versus decay in *Chrysoperla externa* (Ce) for the six best thresholds (overlap length_%identity). Time is controlled for 0, 3 and 6 h after predation of *M. persicae* by *C. externa* for the indirect/ secondary decay in *Ha. axyridis*, and for 0, 3 and 6 h after predation of *C. externa* by *Ha. axyridis* for direct/primary decay in *C. externa*.

| Threshold | | *df* | *SS* | *MS* | *F* | *p* |
|---|---|---|---|---|---|---|
| 100_100 | Ha vs Ce | 1 | 0.17989 | 0.17989 | 2.07 | 0.20981 |
| | Time | 2 | 0.25244 | 0.12622 | 1.45 | 0.31828 |
| | Interaction | 2 | 0.02069 | 0.01035 | 0.12 | 0.89022 |
| | Error | 5 | 0.43465 | 0.08693 | | |
| 100_99 | Ha vs Ce | 1 | 0.19045 | 0.19045 | 2.01 | 0.21541 |
| | Time | 2 | 0.26240 | 0.13120 | 1.39 | 0.33217 |
| | Interaction | 2 | 0.02014 | 0.01007 | 0.11 | 0.90113 |
| | Error | 5 | 0.47362 | 0.09472 | | |
| 125_100 | Ha vs Ce | 1 | 0.22011 | 0.22011 | 2.32 | 0.18805 |
| | Time | 2 | 0.32557 | 0.16278 | 1.72 | 0.27055 |
| | Interaction | 2 | 0.02990 | 0.01495 | 0.16 | 0.85817 |
| | Error | 5 | 0.47394 | 0.09479 | | |
| 150_100 | Ha vs Ce | 1 | 0.27424 | 0.27424 | 3.56 | 0.11776 |
| | Time | 2 | 0.26595 | 0.13297 | 1.73 | 0.26899 |
| | Interaction | 2 | 0.01654 | 0.00827 | 0.11 | 0.90019 |
| | Error | 5 | 0.38495 | 0.07699 | | |
| 150_99 | Ha vs Ce | 1 | 0.31210 | 0.31210 | 4.17 | 0.09654 |
| | Time | 2 | 0.27007 | 0.13504 | 1.81 | 0.25694 |
| | Interaction | 2 | 0.02938 | 0.01469 | 0.20 | 0.82776 |
| | Error | 5 | 0.37399 | 0.07480 | | |
| 175_99 | Ha vs Ce | 1 | 0.26232 | 0.26232 | 3.52 | 0.11932 |
| | Time | 2 | 0.35721 | 0.17861 | 2.40 | 0.18602 |
| | Interaction | 2 | 0.09039 | 0.04520 | 0.61 | 0.58072 |
| | Error | 5 | 0.37224 | 0.07445 | | |

*4) Decay of* C. externa *reads in the predator,* Ha. axyridis. The decay of *C. externa* reads after ingestion by the predator, *Ha. axyridis*, followed a first order decay process and the results were similar for all six thresholds (Fig. S4). There were no significant differences in the decay rate related to the feeding history of *C. externa* (Table S7.1), although it was nearly significant for the 100_100 and 175_99 thresholds. There was no significant correlation between detection of *M. persicae* and *C. externa* reads in *Ha. axyridis* (Fig. S5), implying that detection of each species was independent of the other.

**Table S7.1.** ANOVA of decay rate of *Chrysoperla externa* in *Harmonia axyridis* for different *Chrysoperla externa* feeding histories for the best six thresholds (overlap length_%identity).

| **Threshold** | | ***df*** | ***SS*** | ***MS*** | ***F*** | ***p*** |
|---|---|---|---|---|---|---|
| 100_100 | Feeding history | 3 | 0.2965 | 0.0988 | 9.00 | 0.0521 |
| | Error | 3 | 0.0330 | 0.0110 | | |
| 100_99 | Feeding history | 3 | 0.1004 | 0.0335 | 2.32 | 0.2533 |
| | Error | 3 | 0.0432 | 0.0144 | | |
| 125_100 | Feeding history | 3 | 0.0921 | 0.0307 | 3.01 | 0.1949 |
| | Error | 3 | 0.0306 | 0.0102 | | |
| 150_100 | Feeding history | 3 | 0.0803 | 0.0268 | 2.93 | 0.2003 |
| | Error | 3 | 0.0274 | 0.0091 | | |
| 150_99 | Feeding history | 3 | 0.0728 | 0.0243 | 4.29 | 0.1313 |
| | Error | 3 | 0.0170 | 0.0057 | | |
| 175_99 | Feeding history | 3 | 0.0732 | 0.0244 | 7.54 | 0.0656 |
| | Error | 3 | 0.0097 | 0.0032 | | |

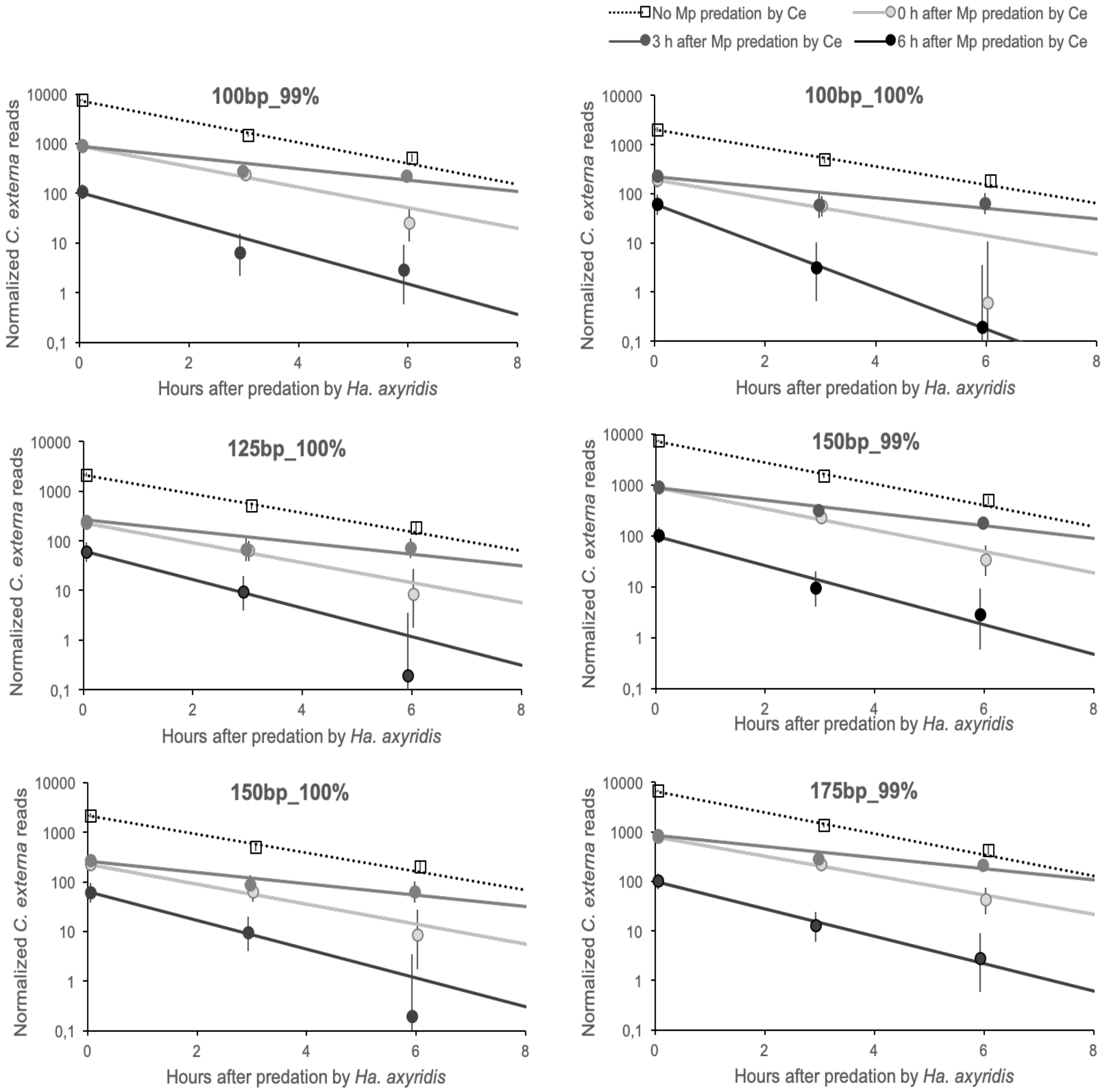


**Figure S4.** Direct predation. Decay of *Chrysoperla externa* reads in *Harmonia axyridis* for the six best thresholds (overlap length_%identity).

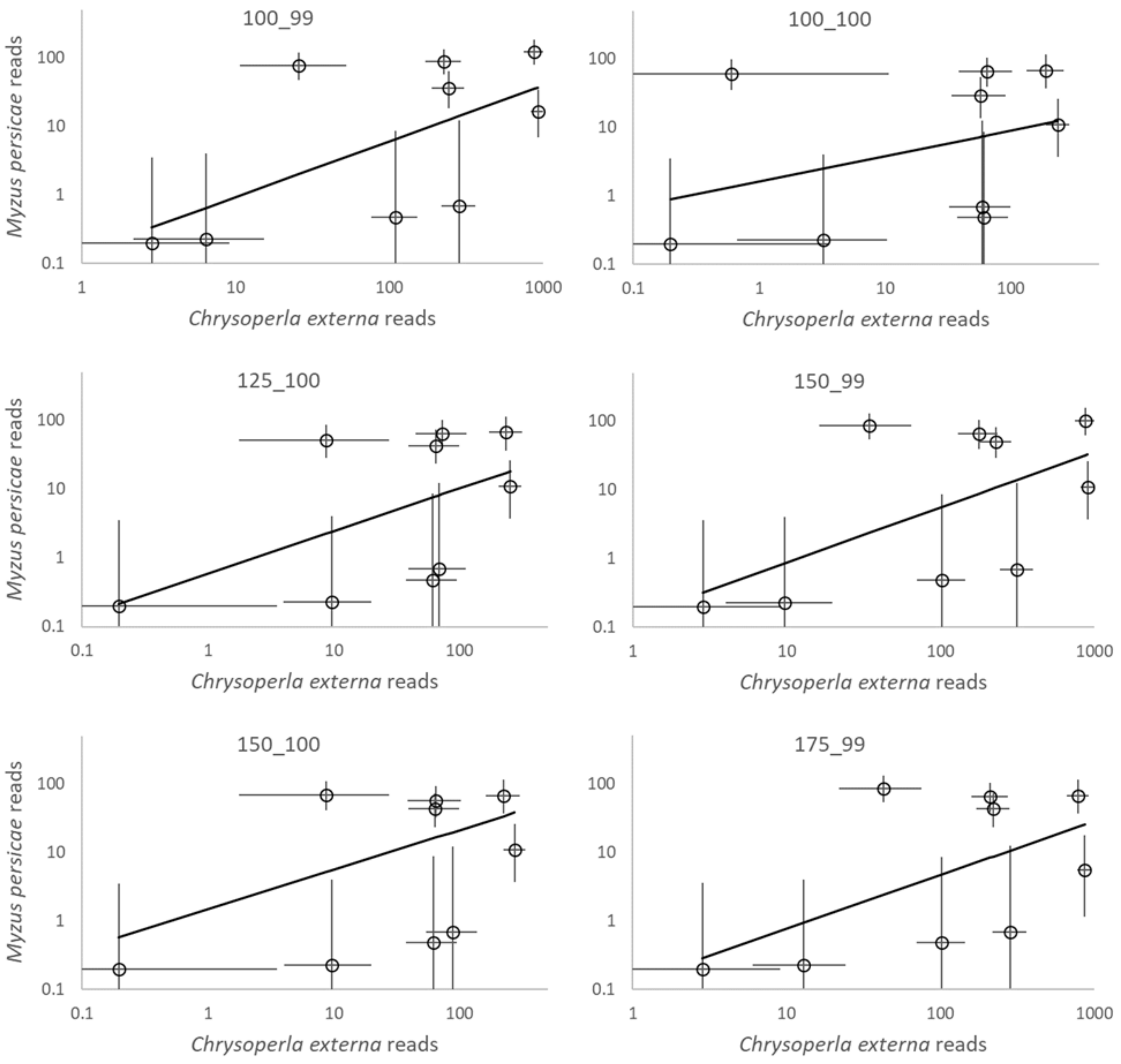


**Figure S5.** Relation between the number of *Myzus persicae* and *Chrysoperla externa* reads detected in *Harmonia axyridis* for the six best thresholds (overlap length_%identity). Pearson correlation coefficients varied from 0.373 to 0.610 ($p$ = 0.503 to 0.177).

**False positive cleaning**

False positive cleaning was didactically divided in two parts: A) reads with single species hits and B) reads with multiple species hits.

A) Reads with single species hits: Reads with single species hits were classified as hits to the predator, hits to the prey and hits to any other species (false positives). There were 174 false positive species in the libraries after BlastNToSnp. In addition, we examined how the thresholds reduced different kinds of false positive reads. These were false positives that were potential cross-contaminant species (contaminants), mismatched species that were in the same family as any of the experimental species (mismatch), and all other species (Other). Potential cross-contaminant species were ones that were in the described experiments or in experiments that were conducted simultaneously. The experimental species were *Hi. convergens*, *Ha. axyridis*, *M. persicae*, *C. externa*, *Cycloneda sanguinea*, and *Ephestia kuehniella*. The first four species were used in the described experiments, *Cy. sanguinea* was a coccinellid predator used in another experiment, and *E. kuehniella* was used as a supplemental food for the predator larvae prior to the experiments. Thus, some of the false positive potential cross-contaminant reads could be due to contamination or carryover of rearing foods in predator guts. Mismatch species are ones that are in the same family as one of the experimental species. There were originally 53 mismatch species. No mismatches to *E. kueniella* were observed. Mismatches may occur from sequencing errors, genetic variation not adequately covered in the reference database or contamination from species not in the experiment but common in the local environment. “Other” species are ones that should not have been detected and could be called “true false positives.” These might occur because the reads are associated with hypervariable regions and minor sequencing errors result in an erroneous determination. There were originally 116 other species.

We estimated the percent reduction in these three kinds of false positives for all 25 thresholds after pairing. In general, the percent removal of reads and detected species of the potential cross-contaminant false positive species was highest only when there was both high overlap length (≥175 bp) and high percent identity (100%). Percent removal of mismatch false positive reads for the *Hi. convergens* libraries decreased with higher overlap length for lower percent identities not because there were more reads retained, but because the original number of mismatched false positive reads was much lower for these cases. Otherwise the thresholds removed 50-90% of the mismatch reads and 70-95% of the mismatch species. Nearly all of the true false positives were removed at ≥99% identity for all of the overlap lengths.

All the six potential cross-contaminant species remained after pairing reads for all of the thresholds. Mismatch species after pairing reads from the thresholds were reduced to 22 species: the 11 aphids *Acyrthosiphon pisum*, *Aphis citricidus*, *A. coreopsidis*, *A. gossypii*, *A. spiracoela*, *Hormaphis betulae*, *Paracolopha morrisoni*, *Schizaphis scirpi*, *Sitobion avenae*, *Therioaphis trifolii* and *Uroleucon ambrosiae* (Hemiptera: Aphididae), the 8 ladybirds *Adalia bipunctata*, *Anisosticta novemdecimpunctata*, *Calvia decemguttata*, *Coccidula rufa*, *Ha. quadripunctata*, *Hi. tredecimpunctata*, *Hi. undecimnotata* and *Propylea japonica* (Coleoptera: Coccinellidae, Coccinellinae), and the three lacewings *Chrysopa pallens*, *Chrysoperla nipponensis* and *Nothochrysa* sp. (Neuroptera: Chrysopidae). The true false positives species after pairing were reduced to nine species: *Amara communis* and *Liogluta microptera* (Coleoptera), *Hydrotaea leucostoma* (Diptera), *Aphidius gifuensis*, *Bombus pascuorum* and *Macrocentrus camphoraphilus* (Hymenoptera), *Helicoverpa zea* (Lepidoptera), *Microchorista philpotti* (Mecoptera) and *Protohermes concolorus* (Megaloptera).

The number of reads and number of hit species were counted for each of the 25 thresholds after pairing (five overlap lengths 100, 125, 150, 175, and 200 bp and five percent identities 96, 97, 98, 99 and 100%). We looked for the thresholds that retained true positive predator or prey reads while eliminating false positive reads and retained the prey species while eliminating false positive species (Fig. S6) according to the three criteria mentioned above. We counted the number of times a given threshold met one of the criteria, and retained the six thresholds that met at least four of the possible six criteria as acceptable thresholds (Table S8). Four thresholds, 100_99 (100 bp overlap length and 99% identity), 125_100, 150_100 and 175_99 met four of the criteria, and two thresholds, 100_100 and 150_99 met five of the six criteria. In all cases the 150_99 threshold eliminated more false positive reads and species than the 100_100 threshold, while retaining the same true positive prey species.

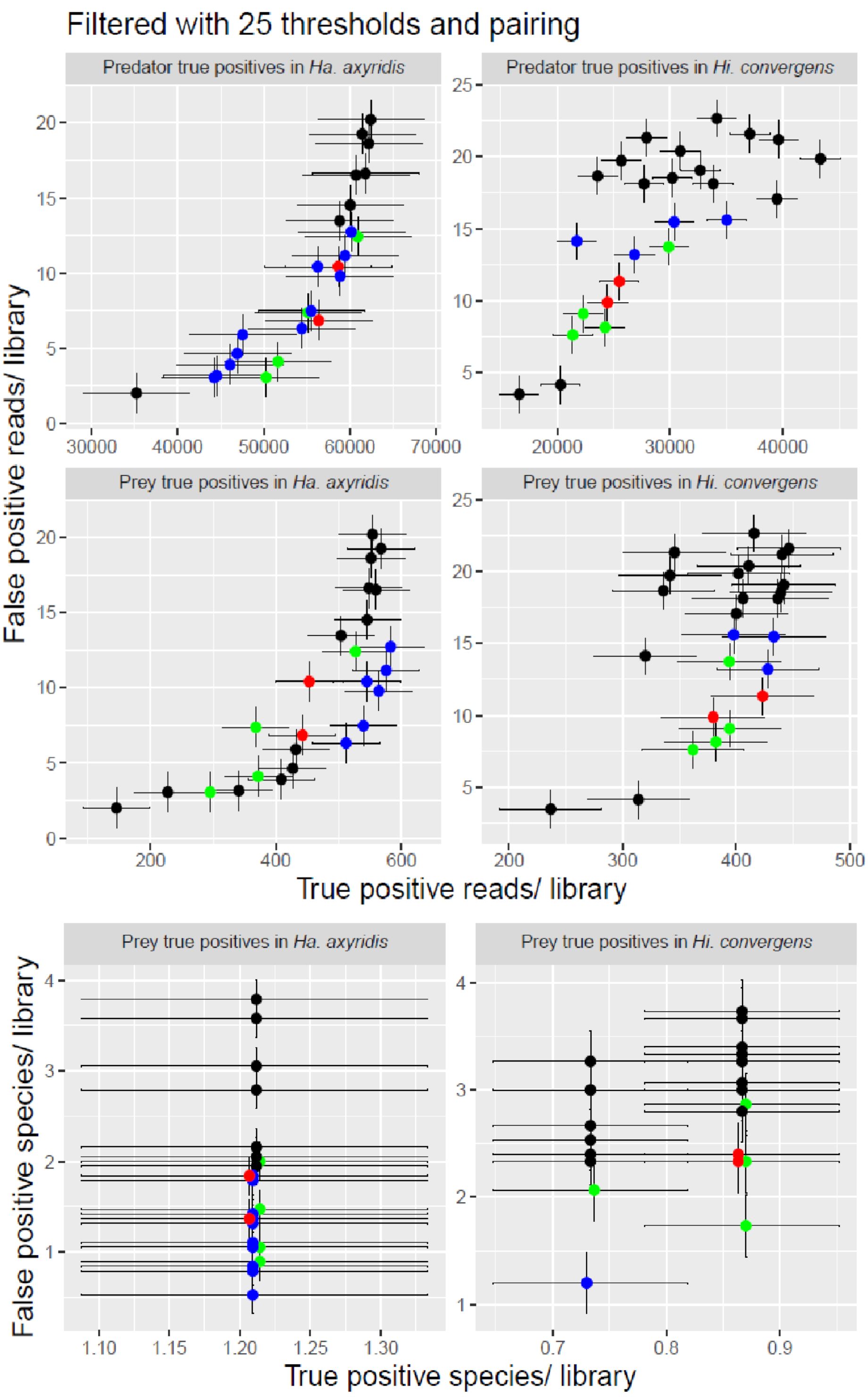


**Figure S6.** Number of false positive reads or species versus true positive reads (predator or prey) or species (prey) for the 25 thresholds in the two experiments with different predators. Black dots are thresholds not meeting any criterion; blue dots are thresholds meeting at least one criterion; green dots are thresholds that meeting 4/6 criteria; red dots are thresholds meeting 5/6 criteria.

**Table S8.** Number of times thresholds met selection criteria.

| Overlap length (bp) | Identity (%) | Number of criteria met |
|---|---|---|
| 100 | 96 | 0 |
| 100 | 97 | 0 |
| 100 | 98 | 2 |
| 100 | 99 | 4 |
| 100 | 100 | 5 |
| 125 | 96 | 0 |
| 125 | 97 | 0 |
| 125 | 98 | 3 |
| 125 | 99 | 3 |
| 125 | 100 | 4 |
| 150 | 96 | 2 |
| 150 | 97 | 2 |
| 150 | 98 | 3 |
| 150 | 99 | 5 |
| 150 | 100 | 4 |
| 175 | 96 | 2 |
| 175 | 97 | 3 |
| 175 | 98 | 3 |
| 175 | 99 | 4 |
| 175 | 100 | 3 |
| 200 | 96 | 2 |
| 200 | 97 | 2 |
| 200 | 98 | 3 |
| 200 | 99 | 3 |
| 200 | 100 | 2 |

The false positive species remaining in any of the six best thresholds after pairing (100_99, 100_100, 125_100, 150_99, 150_100, 175_99) were all the potential cross-contaminant species, eight mismatch species (*Ac. pisum*, *A. coreopsidis*, *A. gossypii* and *U. ambrosiae*, *Ad. bipunctata*, *Ca. decemguttata* and *P. japonica* and *C. nipponensis*), and only one "other" species *Ap. gifuensis*.

Further downstream analysis for the six best thresholds filtered for paired reads that mapped to coding regions of their respective mitogenomes, and lastly subtracted the control library. These analyses greatly reduced the potential cross-contaminant and mismatch false positive reads and species, while retaining a high proportion of true positive prey reads (Fig. S7). All of the potential cross-contaminant species were still detected, although at greatly reduced numbers in all of the thresholds. The most common experimental species was *E. kuehniella* (66% of the reads), the rearing food. The only mismatch species remaining were the aphids, *A. gossypii* and *U. ambrosiae*, the chrysopid, *Ch. nipponensis*, and the coccinellid *Ad. bipunctata*. The only other species was the braconid, *Ap. gifuensis*, but this was detected in only one library in the 100_99 threshold (2 reads total).

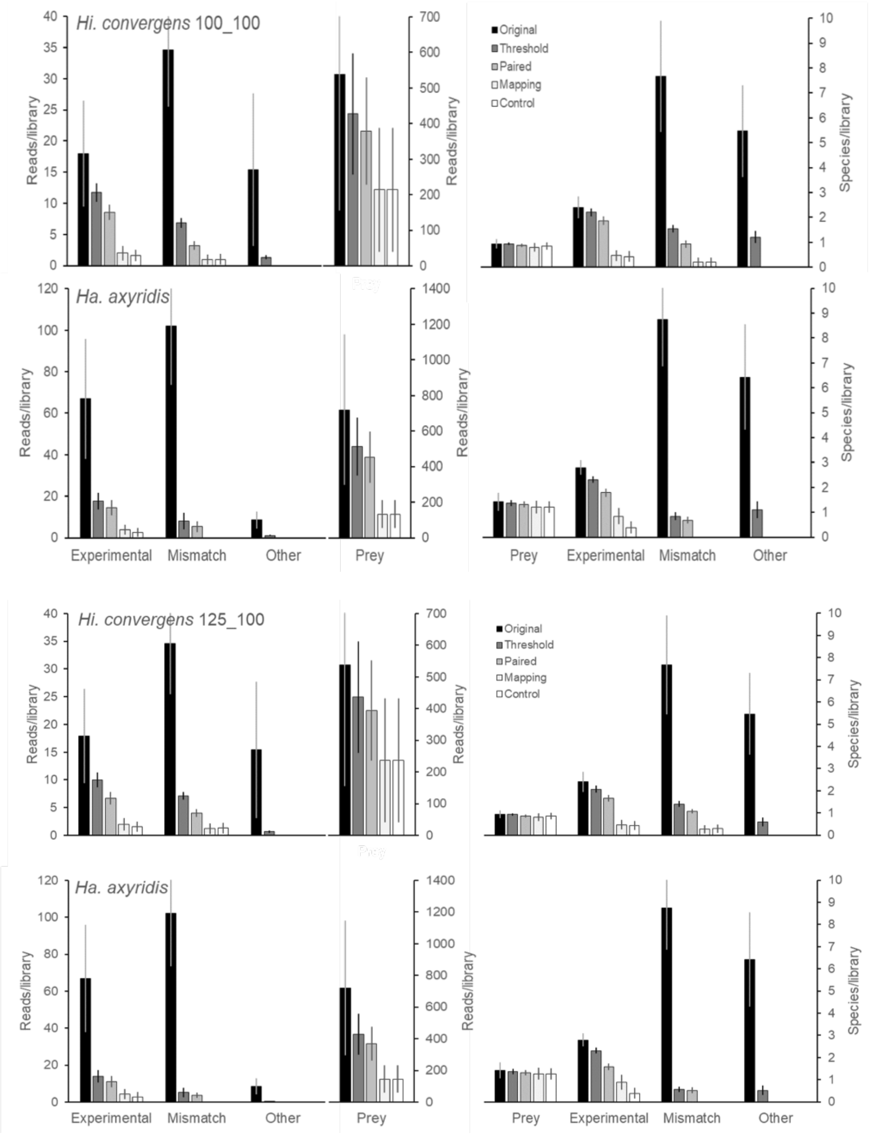

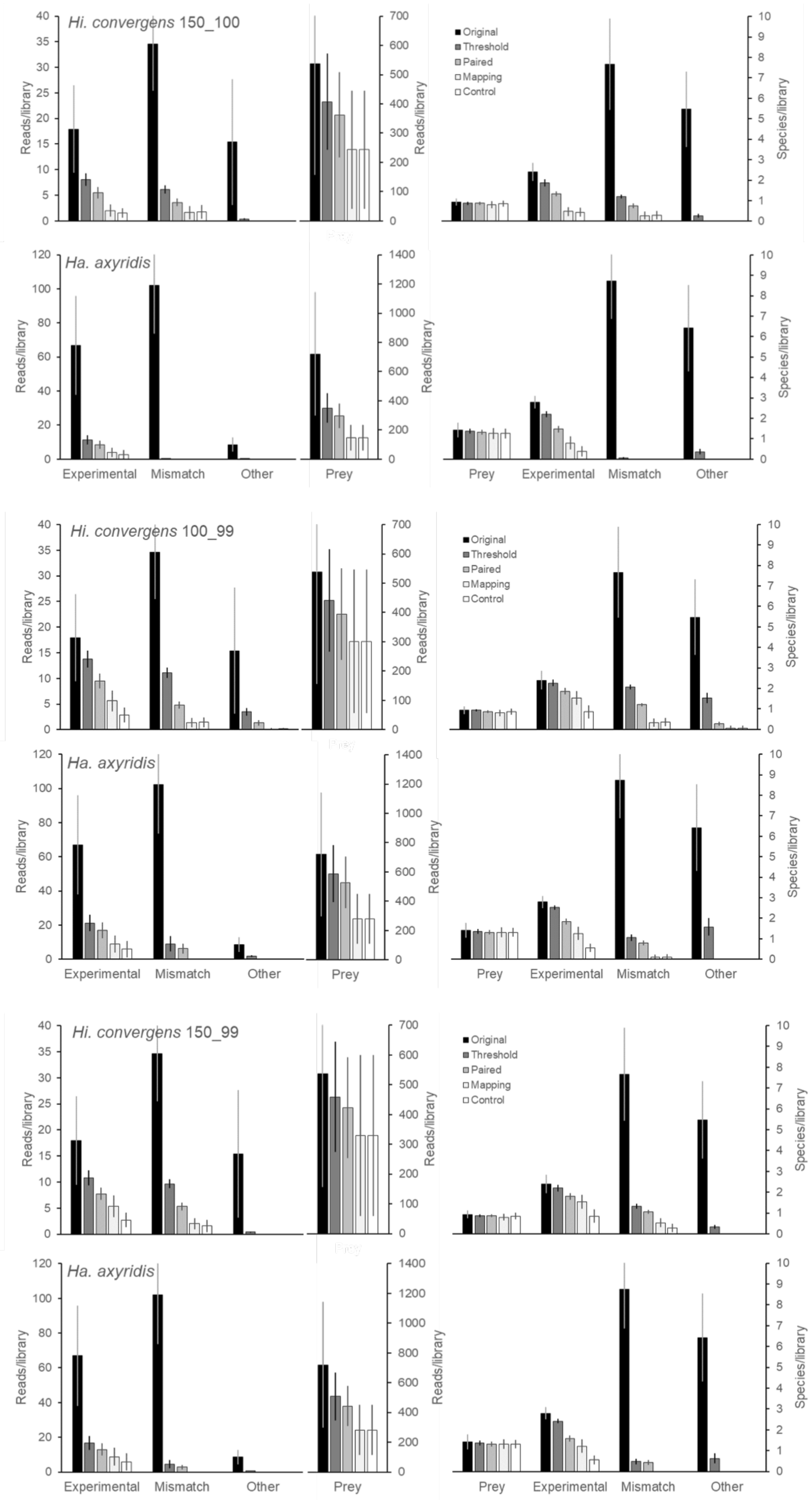
Hi. convergens 150_100
Ha. axyridis
Hi. convergens 100_99
Ha. axyridis
Hi. convergens 150_99
Ha. axyridis
Reads/library
Species/library
Original
Threshold
Paired
Mapping
Control
Experimental
Mismatch
Other
Prey

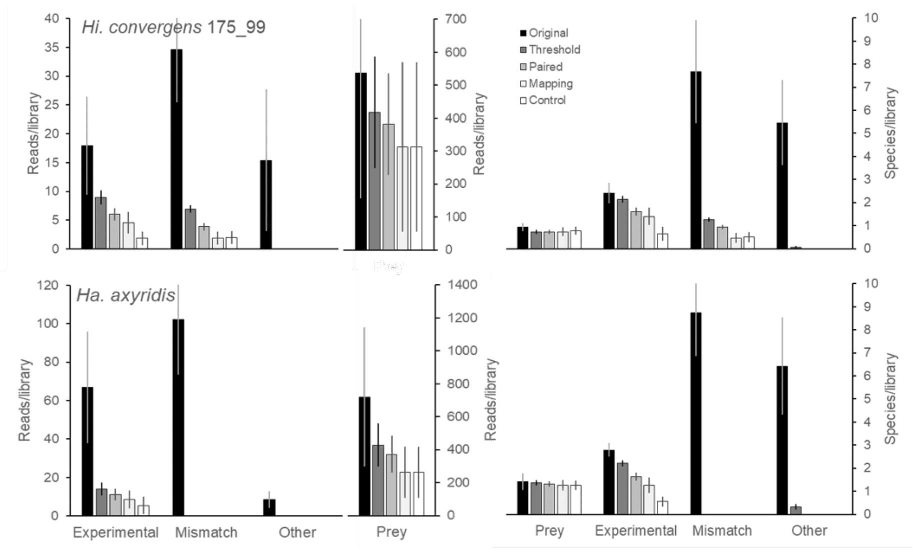


**Fig. S7** Cleaning of false positive reads and species in all the libraries by each of the six best thresholds. Cleaning was conducted by a series of steps: **Threshold:** discard reads with matches below the minimum overlap length and percent identity; **Pairing:** removal of single reads; **Mapping:** removal of reads that do not map with coding regions of the mitogenome of the matches species; **Control:** subtraction of reads that matched with species in the blank library (no-prey consumed by the predator). The false positive potential cross-contaminant reads probably stemmed from DNA contamination of species used in the bioassays. Mismatches are species in the reference database related to ones used in the bioassays and Other are unrelated species. The collateral cleaning effect on the detection of true prey is also shown.

More prey reads were detected with the 99% identity thresholds than the 100% identity thresholds without large increases in false positive reads (Fig. S8). This led us to prefer the 99% identity thresholds over the 100% identity thresholds, because when more prey reads were detected, better estimates of the experimental parameters were possible. Among the 99% identity thresholds, the 100 bp overlap length was the worst at removing false positive species in three cases, the 150 bp overlap length was worst in none of the cases, and the 175 bp overlap length was worst in three cases, so we concluded that the 150bp_99% was the best threshold because it eliminated more false positives than the other 99% identity thresholds. This had four of the six original experimental species (*E. kuehniella*, *Ha. axyridis*, *Hi. convergens* and *Cy. sanguinea*), one of the 53 original mismatch species (*U. ambrosiae*) and no “other” species of 116 original species.

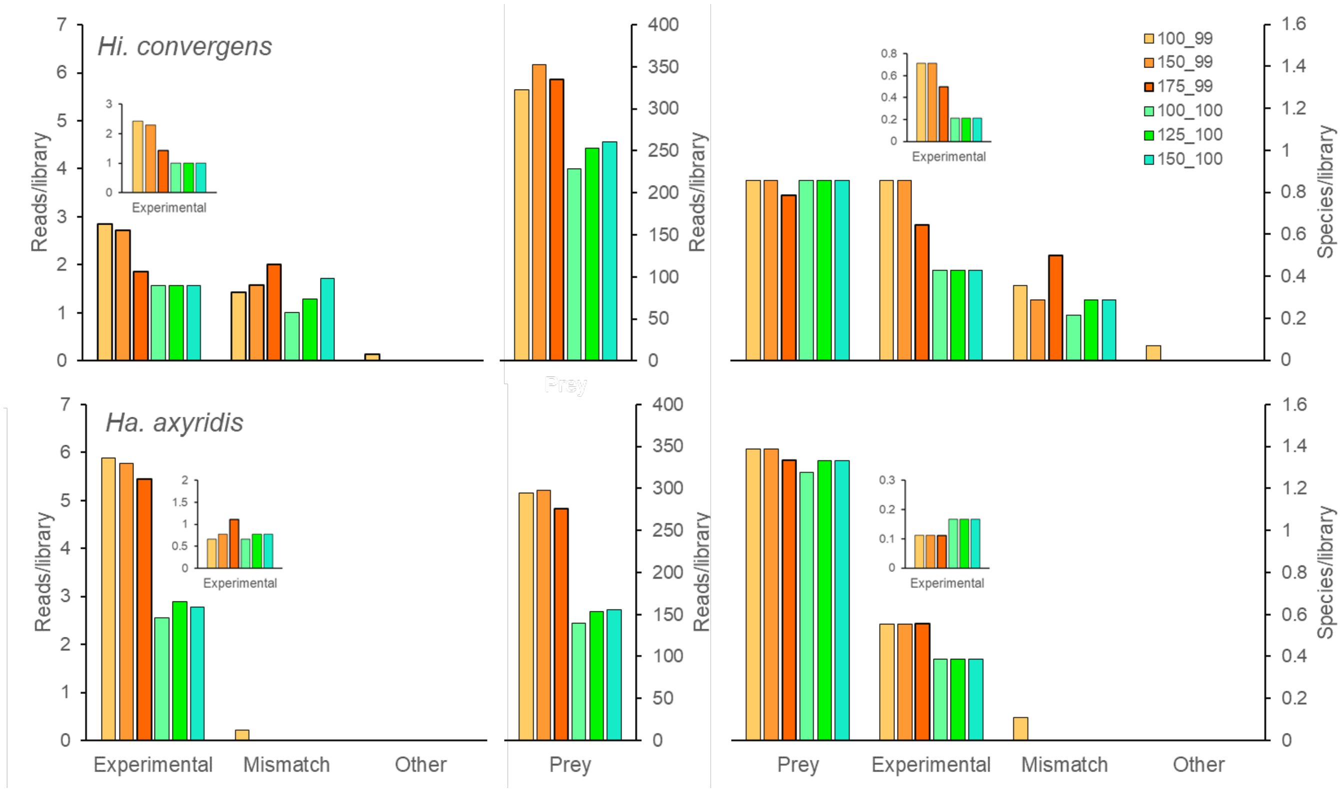


**Fig. S8** Final number of reads/library and species/library after cleaning false positives using the six best thresholds. A) *Hippodamia convergens* and B) *Harmonia axyridis* libraries. Insets are for potential cross-contaminant species without *Ephestia kuehniella*, a rearing food species provided to the predators before the bioassays.

B) Reads with multiple hit species: We eliminated all reads with multiple hits species from the statistical analysis (below) because these reads did not identify a single species. It is likely that many of these reads match to conserved regions of the mitogenome, and therefore do not discriminate species sufficiently to give unique species determinations. However, for field samples, when the true positive species are unknown, the reads with multiple hit species could still be ecologically informative, so we provide analyses to show how the DDSS workflow affected the multiple hit species.

We identified two kinds of reads with multiple hit species. The first kind, and most common in our samples, were reads that hit at least one experimental species. These multi-hit reads provide insufficient discrimination of species, but can provide identifications at higher taxonomic levels. For example, a read with hits on *Hi. convergens* and *Hi. tredecimpunctata* identifies the *Hippodamia* genus, and one that hits *Hi.*

*convergens* and *Ha. axyridis* identifies the tribe Coccinellini. The second kind of reads with multiple hits were reads that did not hit any experimental species. These are true false positives because none of the species hit would be expected under the experimental conditions. We examined how each of these kinds of reads with multiple hits were affected by filtering.

The multiple hits remaining in the best six thresholds after pairing involved aphids, chrysopids and coccinellids (Table S9). There were 119 different multiple hit species combinations involving aphids, of which 91% matched *M. persicae* with an average of 15.9 reads/library, the main prey species in the experiments (Table S9.1A). Of these, 13% matched species in the Macrosiphini, the aphid tribe with *M. persicae*, 64% matches species in the Aphidinae, and the remaining 14% matched species in the Aphididae. The 9% that were *M. persicae* mismatches (aphid matches without *M. persicae*) had only 0.5 reads/library. There were 16 different multiple hit species combinations involving chrysopids, of which 14 matched *C. externa* with an average of 12.1 reads/library (Table S9.1B). These 14 provided a diverse level of taxonomic resolution from genus to superorder. These results suggest that for some taxa, multiple hits provide taxonomic resolution to genus or tribe, but for other taxa the resolution of multiple hits is only to superorder.

**Table S9.** Species included in multiple species hits in the six best thresholds: first group is all species with an Aphididae; second group is species with a Chrysopidae; third group is species with a Coccinellidae; last is a species in the Megaloptera.

| **Species** | **Tribe** | **Subfamily** | **Family** | **Order** |
|---|---|---|---|---|
| *Acyrthosiphon pisum* | Macrosiphini | Aphidinae | Aphididae | Hemiptera |
| *Aphis aurantii* | Aphidini | Aphidinae | Aphididae | Hemiptera |
| *Aphis citricidus* | Aphidini | Aphidinae | Aphididae | Hemiptera |
| *Aphis coreopsidis* | Aphidini | Aphidinae | Aphididae | Hemiptera |
| *Aphis craccivora* | Aphidini | Aphidinae | Aphididae | Hemiptera |
| *Aphis fabae* | Aphidini | Aphidinae | Aphididae | Hemiptera |
| *Aphis glycines* | Aphidini | Aphidinae | Aphididae | Hemiptera |
| *Aphis gossypii* | Aphidini | Aphidinae | Aphididae | Hemiptera |
| *Aphis solanella* | Aphidini | Aphidinae | Aphididae | Hemiptera |
| *Aphis spiracoela* | Aphidini | Aphidinae | Aphididae | Hemiptera |
| *Brevicoryne brassicae* | Macrosiphini | Aphidinae | Aphididae | Hemiptera |

| | | | | |
|---|---|---|---|---|
| *Cavariella salicicola* | Macrosiphini | Aphidinae | Aphididae | Hemiptera |
| *Cinara tujafilina* | Eulachnini | Lachninae | Aphididae | Hemiptera |
| *Diuraphis noxia* | Macrosiphini | Aphidinae | Aphididae | Hemiptera |
| *Eriosoma lanigerum* | Eriosomatini | Eriosomatinae | Aphididae | Hemiptera |
| *Greenidea psidii* | Greenideini | Greenideinae | Aphididae | Hemiptera |
| *Hormaphis betulae* | Hormaphidini | Hormaphidinae | Aphididae | Hemiptera |
| *Hyperomyzus lactucae* | Macrosiphini | Aphidinae | Aphididae | Hemiptera |
| *Kurisakia onigurumii* | Thelaxini | Thelaxinae | Aphididae | Hemiptera |
| *Lipaphis pseudobrassicae* | Macrosiphini | Aphidinae | Aphididae | Hemiptera |
| *Macrosiphum euphorbiae* | Macrosiphini | Aphidinae | Aphididae | Hemiptera |
| *Meitanaphis elongallis* | Fordini | Eriosomatinae | Aphididae | Hemiptera |
| *Mindarus keteleerifoliae* | | Mindarinae | Aphididae | Hemiptera |
| *Myzus persicae* | Macrosiphini | Aphidinae | Aphididae | Hemiptera |
| *Paracolopha morrisoni* | Eriosomatini | Eriosomatinae | Aphididae | Hemiptera |
| *Pterocomma pilosum* | Macrosiphini | Aphidinae | Aphididae | Hemiptera |
| *Rhopalosiphum maidis* | Aphidini | Aphidinae | Aphididae | Hemiptera |
| *Rhopalosiphum padi* | Aphidini | Aphidinae | Aphididae | Hemiptera |
| *Schizaphis graminum* | Aphidini | Aphidinae | Aphididae | Hemiptera |
| *Schizaphis scirpi* | Aphidini | Aphidinae | Aphididae | Hemiptera |
| *Sitobion avenae* | Macrosiphini | Aphidinae | Aphididae | Hemiptera |
| *Uroleucon ambrosiae* | Macrosiphini | Aphidinae | Aphididae | Hemiptera |
| *Abachrysa eureka* | Belonopterygini | Chrysopinae | Chrysopidae | Neuroptera |
| *Chrysopa pallens* | Chrysopini | Chrysopinae | Chrysopidae | Neuroptera |
| *Chrysoperla externa* | Chrysopini | Chrysopinae | Chrysopidae | Neuroptera |
| *Chrysoperla nipponensis* | Chrysopini | Chrysopinae | Chrysopidae | Neuroptera |
| *Eudasyphora canadiana* | | | Muscidae | Diptera |
| *Fontecilla graphicus* | | | Polystoechotidae | Neuroptera |
| *Leucochrysa pretiosa* | Leucochrysini | Chrysopinae | Chrysopidae | Neuroptera |
| *Nipponeurorthus fuscinervis* | | | Nevrorthidae | Neuroptera |
| *Nothochrysa sp.* | | Nothochrysinae | Chrysopidae | Neuroptera |
| *Protochauliodes biconicus* | | | Corydalidae | Megaloptera |
| *Protohermes concolorus* | | | Corydalidae | Megaloptera |
| *Adalia bipunctata* | Coccinellini | Coccinellinae | Coccinellidae | Coleoptera |
| *Aiolocaria hexaspilota* | Coccinellini | Coccinellinae | Coccinellidae | Coleoptera |
| *Amara communis* | | | Carabidae | Coleoptera |
| *Anisosticta novemdecimpunctata* | Coccinellini | Coccinellinae | Coccinellidae | Coleoptera |
| *Calvia decemguttata* | Coccinellini | Coccinellinae | Coccinellidae | Coleoptera |
| *Harmonia axyridis* | Coccinellini | Coccinellinae | Coccinellidae | Coleoptera |
| *Harmonia quadripunctata* | Coccinellini | Coccinellinae | Coccinellidae | Coleoptera |
| *Hippodamia convergens* | Coccinellini | Coccinellinae | Coccinellidae | Coleoptera |
| *Hippodamia tredecimpunctata* | Coccinellini | Coccinellinae | Coccinellidae | Coleoptera |
| *Propylea japonica* | Coccinellini | Coccinellinae | Coccinellidae | Coleoptera |
| *Protohermes concolorus* | | | Corydalidae | Megaloptera |

**Table S9.1.** Taxonomic resolution of multiple species hits (in reads/library) in A) Aphididae, B) Chrysopidae, C) Coccinellidae.

| **A) Taxonomic resolution** | **Match *Myzus persicae*** | **Mismatch** | **Total** |
|---|---|---|---|
| Aphididae | 1.04 | 0.09 | 1.13 |
| Aphidinae | 9.39 | 0.40 | 9.79 |
| Macrosiphini | 5.43 | 0.03 | 5.46 |
| Total | 15.86 | 0.52 | 16.38 |

| **B) Taxonomic resolution** | **Match *Chrysoperla externa*** | **Mismatch** | **Total** |
|---|---|---|---|
| Endopterygota | 0.09 | 0.01 | 0.10 |
| Neuropterida | 0.32 | 0 | 0.32 |
| Neuroptera | 0.03 | 0 | 0.03 |
| Chrysopidae | 0.49 | 0.08 | 0.57 |
| Chrysopinae | 0.03 | 0 | 0.03 |
| Chrysopini | 1.90 | 0 | 1.90 |
| *Chrysoperla* | 9.26 | 0 | 9.26 |
| Total | 12.13 | 0.09 | 12.22 |

| **C) Taxonomic resolution** | **Match *Harmonia axyridis*** | **Match *Hippodamia convergens*** | **Total** |
|---|---|---|---|
| Endopterygota | 0.01 | 0 | 0.01 |
| Coleoptera | 0.02 | 0 | 0.02 |
| Coccinellini | 0.57 | 4925.42 | 4925.99 |
| Genus | 0.01 | 46.01 | 46.02 |
| Total | 0.61 | 4971.43 | 4972.04 |

For the best six thresholds, all “other” multiple hits (did not hit an experimental species or mismatches) and all predator mismatches were filtered out by all of the thresholds. Multiple hit prey mismatches were reduced to ~ 1 read/library in the *Hi. convergens* libraries. In the *Ha. axyridis* libraries, the thresholds with overlap of 100 bp had ~1 read/library, and the other four thresholds had ~0.1-0.2 prey mismatches/library. This suggests that the thresholds with 100 bp overlap did not filter the multiple hit prey mismatches as well as the other thresholds.

C) KrakenUniq: Although we did not measure processing speed, KrakenUniq was considerably faster than the DDSS workflow. The DDSS workflow typically found a different number of reads than KrakenUniq (Table S10). In one assay, DDSS found more predator reads than KrakenUniq, but in the other assay the result was reversed. In one assay the number of prey reads was similar for the two methods, but in the other assay, KrakenUniq found more prey reads. The only consistent result was that KrakenUniq found more false positive reads. The false positive reads from KrakenUniq included one "true" false positive species, *Ap. gifuensis*, a species that is unrelated to any of the experimental species, which was not detected by DDSS. The two methods found similar number of mismatch species reads, but KrakenUniq found ~8 times more false positive reads of potential cross-contaminant species. These false positive potential cross-contaminant reads were only 13% *E. kuhniella* (a larval rearing food) for KrakenUniq, while they were 65% *E. kuhniella* for DDSS. Thus, DDSS was more efficient at removing the possible contamination with experimental species. It is not possible to conclude that one of these methods was more sensitive at finding true positive reads, but it is clear that KrakenUniq found more false positives. This resulted in a positive predictive value (PPV) of 90.5% for KrakenUniq versus 98.4% for the DDSS workflow. Additional experimental evidence will be needed to determine which method is more appropriate for gut contents analysis, but in this experiment, the DDSS workflow was superior.

**Table S10.** Average number of reads per library for KrakenUniq and DDSS with a test of the null hypothesis that the numbers are equal for both methods using a paired *t*-test with libraries as replicates (*n*). Differences were transformed to be normal. Library means and standard errors for each method. FP= false positive.

| Assay | Species | Role | KrakenUniq | DDSS | *n* | *t* | *p* |
|---|---|---|---|---|---|---|---|
| Prey quantity | *Hi. convergens* | Predator | 44435 (2184) | 19864 (1228) | 14 | 9.624 | 2.80E-7 |
| Prey quantity | *M. persicae* | Prey | 363 (189) | 353 (182) | 13 | 0.874 | 0.3995 |
| Prey quantity | *C. sanguinea* | FP | 93.3 (30.3) | 1.25 (0.37) | 8 | 3.012 | 0.0196 |
| Prey quantity | All others | FP | 11.38 (1.58) | 2.08 (0.6) | 24 | 5.501 | 1.36E-5 |
| Indirect | *Ha. axyridis* | Predator | 33795 (2734) | 53727 (4312) | 18 | -11.161 | 3.03E-9 |
| Indirect | *C. externa* | Prey | 342 (178) | 229 (126) | 12 | 9.905 | 8.14E-7 |
| Indirect | *M. persicae* | Prey | 203 (88) | 173 (81) | 11 | 3.32 | 0.0077 |
| Indirect | All | FP | 14.39 (3.94) | 6.11 (2.77) | 17 | 3.728 | 0.0018 |
